# Single-library chromosome-scale diploid assemblies of vole genomes resolve a species-specific duplication implicated in pair bonding

**DOI:** 10.64898/2026.03.13.711624

**Authors:** Mohamed Abuelanin, Gulhan Kaya, Juniper A. Lake, Christine Lambert, Melody V. Wu, Kristen M. Berendzen, Jonathan M.D. Wood, Ksenia Krasheninnikova, Nancy G. Solomon, Zoe R. Donaldson, Karen L. Bales, Kerstin Howe, Jonas Korlach, Devanand Manoli, Jessica Tollkuhn, Megan Y. Dennis

## Abstract

Comparing prairie voles (*Microtus ochrogaster*), a species that forms lasting pair bonds and exhibits biparental care, with meadow voles (*Microtus pennsylvanicus*), a non-monogamous species showing maternal care only, provides a framework to examine the genomic basis of behavioral divergence over short evolutionary time. Here, we develop a simplified assembly approach combining PacBio HiFi and CiFi sequencing in a single library and run, producing high-quality contiguous chromosome-scale diploid prairie and meadow vole genomes (2.3 Gbp). Comparative analysis reveals substantial differences, including near-complete Y chromosome divergence and a prairie-vole-specific duplication of *Avpr1a*, a vasopressin receptor gene implicated in social bonding and autism in humans. These extensive genomic changes suggest rapid chromosome evolution as a driver of the dramatic *Microtus* radiation, generating ∼60 vole species in >2 million years. This single-library approach facilitates a simplified and more affordable assembly workflow, producing near-complete genomes of diverse species using one sequencing platform.

## Introduction

High-quality reference genomes are foundational for characterizing the genomic basis of speciation, phenotypic diversity, and disease susceptibility^1^. They provide the substrate to identify functional genes and regulatory regions, and to overlay variants that impact them. However, repetitive and structurally complex regions, including segmental duplications (SDs) and centromeres, have historically been underrepresented or absent from reference assemblies. These regions are recognized as a primary source of new gene functions for adaptive evolution^2^, are vital for basic cellular functions (centromeres), and, when mutated, can lead to human disease. Advances in long-read sequencing, particularly PacBio HiFi technology, have transformed *de novo* genome assembly by producing highly accurate reads exceeding 10 kbp^3^. When combined with chromosome conformation capture (3C) data, chromosome-scale, fully phased, haplotype-resolved assemblies can be generated^4^. Genome-wide 3C (Hi-C) depends on short-read sequencing, which maps poorly to repetitive regions and necessitates multi-platform library preparations^5,6^. CiFi addresses these limitations by integrating 3C libraries with PacBio HiFi long-read sequencing^7^, validated on a Mediterranean fruit fly genome assembly (∼600 Mbp) and requiring lower coverage compared with Hi-C for effective scaffolding. Since both DNA products can be sequenced on a uniform platform and separated bioinformatically based on distinct sizes/adapters, there is an opportunity to combine HiFi/CiFi into one library preparation and optimized workflow reducing expense and technical complexity.

Here, we demonstrate a single-library HiFi-CiFi assembly approach for two mammalian genomes approximately four-fold larger (∼2.3 Gbp) than the previously sequenced fruit fly: prairie vole (*Microtus ochrogaster*) and meadow vole (*Microtus pennsylvanicus*). Voles diverged from the murine lineage, including mice and rats, approximately 20 million years ago (mya)^8^. Mice and rats (within subfamily *Murinae*), both extensively used as model organisms, share a common ancestor estimated at ∼12–23 mya, while the *Arvicolinae* subfamily comprising voles originated ∼6–7 mya, with its most dramatic radiation occurring within the genus *Microtus* ∼2 mya^9,10^. Similar to humans, prairie voles are among the few mammalian species exhibiting social monogamy^11,12^. They form enduring social attachments to their partners, known as pair bonds^13^, and both parents care for offspring. In contrast, closely related meadow voles do not show social monogamy, and only mothers care for offspring. Thus, the neurogenetic differences between vole species offer a powerful system to examine the mechanistic basis of complex social behaviors. Further, prairie vole behaviors better parallel aspects of human interactions compared with mice or rats, making them a valuable model for understanding the neural basis of human social behaviors. Across taxa, neuropeptide hormones represent a critical substrate in the evolution of social behaviors, modulating the neural circuits and underlying physiology that control social interactions^14,15^. Seminal comparative behavioral and pharmacologic studies have implicated the nonapeptide hormones oxytocin and vasopressin in prairie vole social attachment^16–19^. Oxytocin receptor (*Oxtr*) and vasopressin receptor 1a (*Avpr1a*) exhibit divergent brain expression patterns in prairie voles relative to other vole species, suggesting substantial regulatory divergence at these loci^20,21^.

The existing prairie vole reference (MicOch1.0; Broad Institute, 2012) predates modern long-read technologies—generated from a 94× coverage Illumina sequence of a female individual—and lacks chromosome-scale contiguity and haplotype resolution. Notably, a >105 kbp prairie-vole duplication of *Avpr1a*, identified by targeted BAC sequencing^22^, was not present in MicOch1.0. Subsequent work on the *Avpr1a* paralog has been hampered, including verification of its existence in prairie vole or other related species, due to the complex nature of this locus. A chromosome-scale meadow vole assembly was recently generated by the Vertebrate Genomes Project (VGP mMicPen1^1^) but has not been compared to the prairie vole in the context of social behavior genetics. Using a new workflow that combines HiFi and CiFi DNA for a single library preparation and sequencing run per vole species, we produce haplotype-resolved, chromosome-scale references enabling a systematic assessment of species divergence.

## Results

### HiFi-CiFi single-library sequencing and genome assembly

To generate HiFi and CiFi data from a single PacBio sequencing library, we isolated high-molecular-weight genomic DNA (gDNA) and fixed chromatin from liver tissue of an individual male prairie vole and meadow vole, respectively (Figure 1A). HiFi gDNA was sheared to a target average fragment size of 16 kbp, while CiFi DNA was prepared using the restriction enzyme HindIII and standard protocol^7^, yielding average fragment sizes of ∼8 kbp (Figure S1). The two preparations were pooled at a 90:10 molar ratio (HiFi:CiFi), constructed into a single library, and sequenced on one Revio SMRT Cell per species, producing 102.7 Gbp (prairie vole) and 98.2 Gbp (meadow vole) of data at a median read quality of QV38.

**Figure 1.**
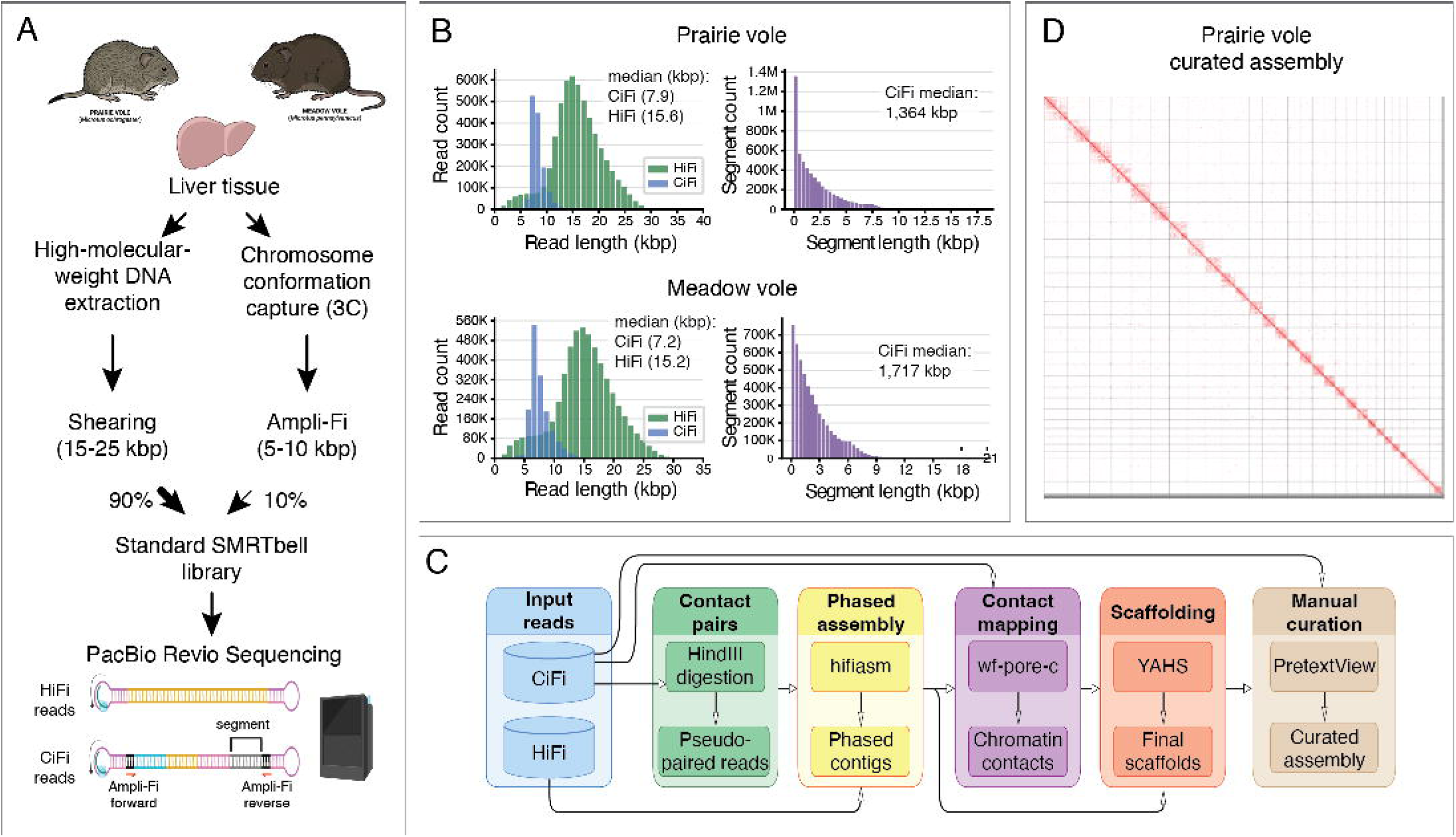
Prairie and meadow vole single-library HiFi-CiFi workflow and sequencing statistics. **(A)** Library preparation from liver tissue using DNA for HiFi (left) and chromatin for CiFi (right) combined (90:10) to construct a single SMRTbell library. **(B)** Read-length (HiFi and CiFi) and segment-length (CiFi) distributions. **(C)** Computational pipeline for HiFi-CiFi-based genome assembly and scaffolding. **(D)** CiFi contact map of the curated prairie vole Hap1 assembly. Uncurated contact maps for both haplotypes and species are in Figures S2 and S3. See also Figures S1 and S4–S7, Tables S1–S4, and Note S1.

HiFi (88%) and CiFi (12%) reads were segregated using unique Ampli-Fi adapter sequences flanking CiFi fragments (Methods, Figure 1A). HiFi reads showed median lengths of 15.6 kbp (prairie vole) and 15.2 kbp (meadow vole) at 35–40× predicted coverage, while CiFi reads were shorter, as expected, producing 5× predicted coverage (Figure 1B, Table S1). CiFi reads were then converted from multi-segment concatemers into chromatin-contact pairs: 1.37 million CiFi reads yielded 23.4 million contact pairs for prairie vole, and 1.54 million reads yielded 8.1 million pairs for meadow vole. The greater contact-pair yield for prairie vole reflects slightly smaller segment sizes (median 1.4 vs 1.7 kbp) and longer CiFi read lengths (median 7.9 vs 7.2 kbp). Together, these results demonstrate that a combined HiFi and CiFi library produces high-quality data.

In order to assess CiFi as a single-platform alternative to Hi-C for *de novo* genome assembly, we generated parallel short-read Hi-C data from the same liver tissues using the standard DpnII restriction enzyme (∼49 Gbp, 20× for prairie vole; ∼73 Gbp, 30× for meadow vole). HiFi reads with either CiFi-derived contact pairs or Hi-C read pairs were used to generate phased diploid contig assemblies with hifiasm^4^, then scaffolded with YAHS^23^ (Figure 1C). CiFi-phased contigs were comparable or longer in three of four haplotypes and achieved better or equivalent scaffold consolidations, approaching expected chromosome numbers more closely than Hi-C (Figures S2–S4, Table S2, Note S1). Downsampling CiFi reads, we observed near-optimal scaffold N50 of all four haplotypes at 40% input or 5 Gbp of data (∼2× genome coverage) (Figure S5). We next extended this test to a human genome (3.2 Gbp) where 2× CiFi and 30× HiFi are achievable on a single SMRT Cell. Comparing a hifiasm assembly generated with 2× CiFi^7^ against pedigree-phased NA12878 variants^24^, we observe highly accurate haplotype phasing: median per-chromosome switch error was 0.18% versus 0.38% for HiFi only, and essentially equal to standard 30× Hi-C (0.16%) (Figure S6, Table S3, Methods S1). Together, this indicates that 2× CiFi is suitable to achieve equal to better phasing and contiguity assembly metrics compared to a standard ∼30× Hi-C.

### Final curated vole assemblies

Assemblies produced with HiFi and CiFi required minimal correction during manual curation (see Methods), including five scaffold breaks and four joins for prairie vole and, similarly, five scaffold breaks and ten joins for meadow vole (Figures 1D, S2, S3, and S7). The sex chromosomes were identified for each species (represented in Hap1), with the remaining autosomes named according to descending size. Assessment of curated assemblies show both haplotypes of each diploid vole assembly to be contiguous (contig N50 > 39 Mbp; scaffold N50 > 89 Mbp), complete (genomic BUSCO ≥99%), and high quality^25^ (diploid QV 50.9–51.9), with diploid heterozygosity of 0.99% (prairie vole) and 1.14% (meadow vole) (Tables 1 and S4, Figure S8).

The prairie vole diploid assembly represents a complete karyotype (2n = 54)^26,27^ with 52 autosomes, chrX, and chrY (Total: 4.83 Gbp, Hap1: 2.53 Gbp). Telomeres were detected on all but three chromosomes (mean 3.5 gaps/chr). Of these, 31 represent T2T scaffolds and three gapless contigs (overall 63%) (Figure 2, Table S5). In contrast, the current prairie vole reference (MicOch1.0)—a haploid assembly using short-read sequencing of a female individual—comprises 18 chromosomes (17 autosomes and chrX), 10 linkage groups, and four unlocalized scaffolds (1.66 Gbp). 632 Mbp of sequence is represented as 6,303 unplaced scaffolds versus ∼30 Mbp in 69 unplaced scaffolds in prairie-vole Hap1 from this study. Comparing the assemblies also revealed major improvements in contiguity (e.g., contig N50: 29.2 kbp MicOch1.0 vs. 39.2 Mbp Hap1) and reduced gap content (8.0% vs. 0.07%), with 238 previously unplaced MicOch1.0 scaffolds (520.8 Mbp) now incorporated in Hap1 chromosome-scale scaffolds (Figure S9, Tables S6 and S7).

**Figure 2.**
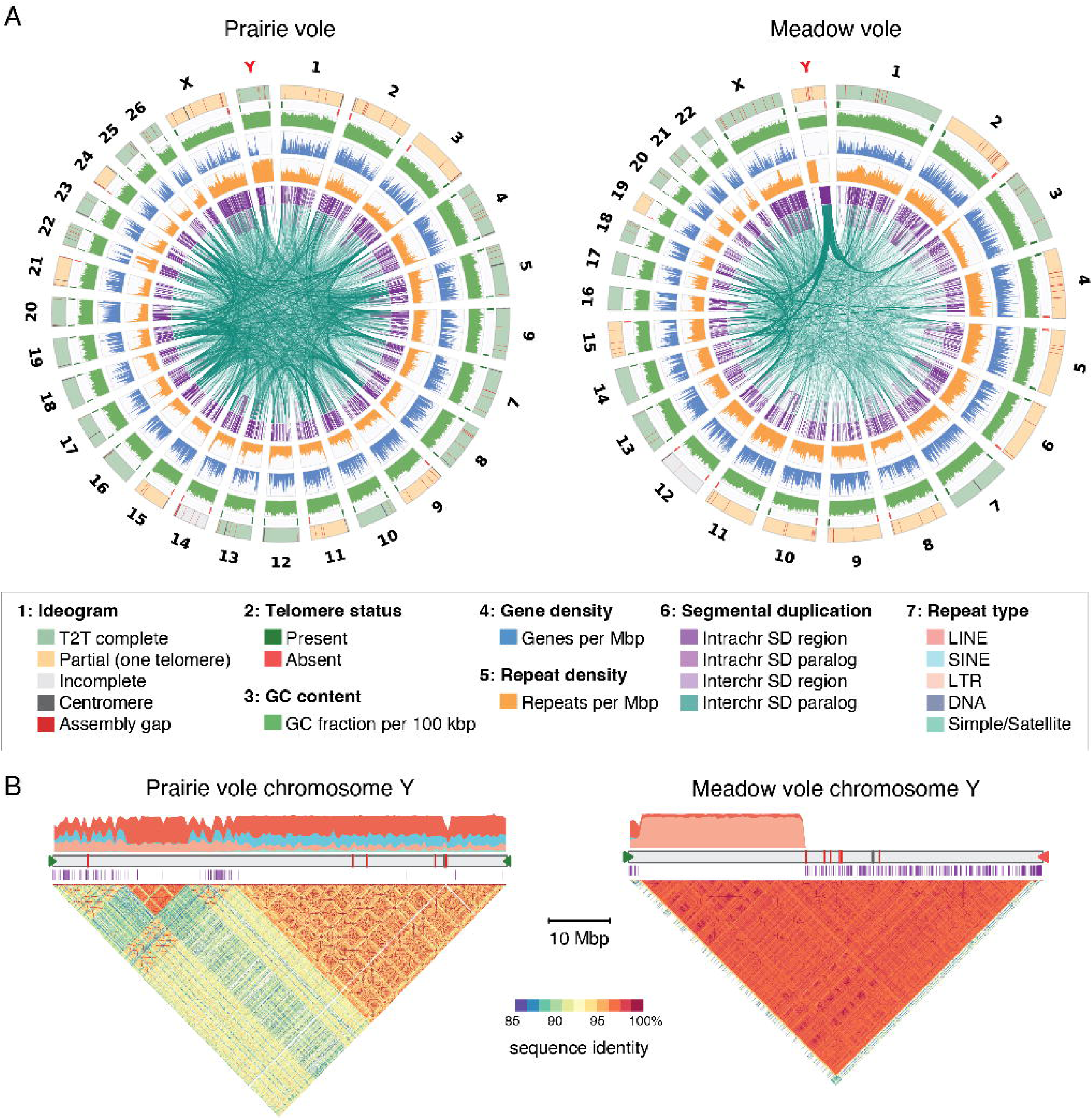
Prairie and meadow vole assembly characteristics. **(A)** Circos plots depicting chromosome characteristics of Hap1assemblies for prairie (left) and meadow (right) vole assemblies. Tracks are described in the legend from outer to inner ring (1: Ideogram; 6: segmental duplication (SD)). **(B)** Repeat and SD distributions are shown above and below the chromosome Y ideogram for each species. Ideogram annotations follow the color scheme in (A), indicating centromere and assembly gap (legend 1), telomere status (legend 2), SDs (purple; legend 6), and repeat types (legend 7). Dotplots below each ideogram show chromosome Y sequence identity. Annotated ideograms of all Hap1 chromosomes are shown in Figures S11 and S12. See also Figure S8–S10 and Tables S8–S10.

Similarly, the meadow vole assembly matches the expected karyotype (2n=46)^28,29^ across 44 autosomes, chrX, and chrY, representing 4.51 Gbp total sequence (Hap1: 2.36 Gbp). All but two chromosomes have at least one telomere detected (mean 3.4 gaps/chr), with 17 T2T scaffolds and six gapless contigs (overall 50%; Figure 2, Table S5). A majority of the meadow vole autosomes are telocentric (42/44), with the two sex chromosomes classified as subtelocentric^28,29^, likely contributing to the higher proportion of single telomeres detected in the meadow versus prairie vole. This assembly is on par with the recent long-read VGP meadow vole reference (mMicPen1), which has all expected chromosomes at near equal contiguity (scaffold Hap1 N50: 125.8 Mbp vs. this study 125.3 Mbp) but a larger number of unanchored sequences (n=156 at 46.1 Mbp vs. this study n=81 at 19.7 Mbp; Figure S10).

Moving forward, we characterized primary Hap1 chromosomes, comprising single sets of autosomes and both sex chromosomes. Using RNA-seq data from liver, brain, and testes of adults and e11.5 (prairie vole) or e13.5 (meadow vole) embryos, we annotated ∼20,500 protein-coding genes per species (Methods, Figure 2, Table S8). Repeat annotation revealed an increased proportion of interspersed repeats in prairie vole (e.g., L1 elements representing 16.7% vs. 13.0% of the genome), satellite repeats (8.62 Mbp vs. 511 kbp), and unclassified repeats (>40 Mbp more than meadow vole) (Figures 2A, S11, S12, Table S9). Near-equal numbers of segmental duplications (SDs; regions >1 kbp at >90% sequence identity^30,31^) were identified in both genomes (∼5.3K), with 72–79% intersecting an annotated gene (prairie vole: 1,151 genes; meadow vole: 1,194 genes). Despite this similarity, prairie vole harbored nearly double the duplicated sequence (54 Mbp, 2.15% of genome) compared to meadow vole (28 Mbp, 1.2%) (Figures 2A, S11, S12, and Table S10). This expanded SD content is driven by larger duplicons (15.2 kbp vs. 8.21 kbp) and a greatly expanded interchromosomal SD fraction (34.7 Mbp, 63.7% of pairs vs. 9.7 Mbp, 46.2%).

We next examined the sex chromosomes given their known rapid and repeated structural remodeling in *Microtus*^32,33^. The most notable differences are evident on the Y chromosome (Figure 2B): both species showed gene depletion and likely constitutive heterochromatin spanning either half (prairie vole) or the entirety (meadow vole) of the chromosome. The remaining half of the prairie vole chrY comprises tandemly arrayed SDs of a *Usp9y* mouse homolog, while the meadow vole chrY also carries considerable SD content. Direct sequence comparison reveals no orthology (Figure 3A), indicating that homologous Y chromosomes can reach complete sequence divergence over remarkably short evolutionary timescales.

**Figure 3.**
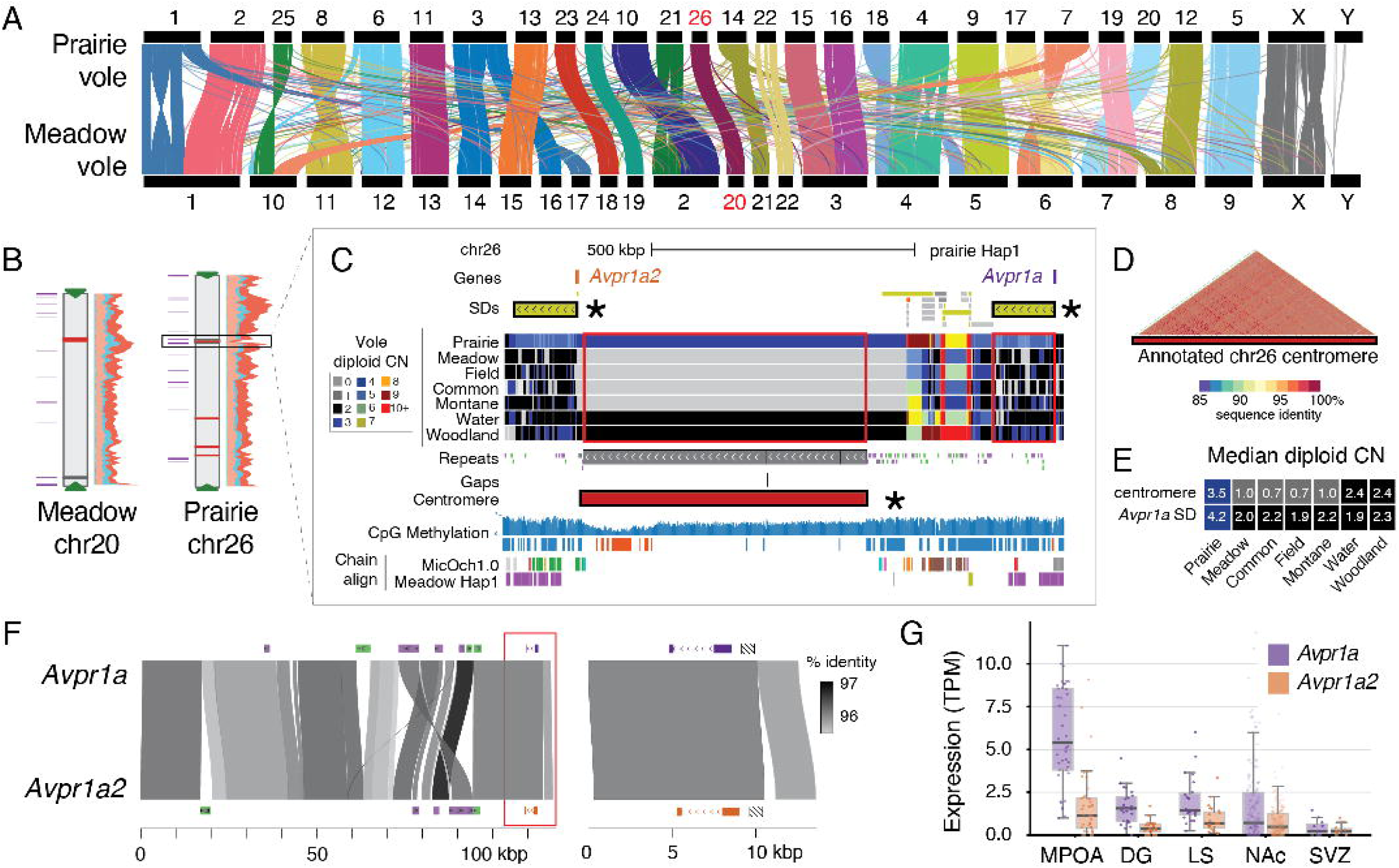
Prairie and meadow vole cross-species comparison. (**A**) Ribbon plots depicting synteny between voles^72^ (colored by homologous chromosome) with chromosome numbers indicated above and below. (**B**) Repeat (right) and segmental duplication (SD; left) annotations of the homologous chromosomes carrying *Avpr1a*. Color scheme as in Figure 2B. (**C**) UCSC Genome Browser view of the prairie vole Hap1 *Avpr1a* locus (chr26:13.0 Mbp–14.1 Mbp). SDs of paralogs and intervening centromeric repeat are highlighted with an asterisk. Windowed copy numbers (CN) of vole species are depicted by colors. CpG methylation is shown as the proportion of methylated HiFi reads (0 to 1); methylated (blue) and hypomethylated (red) regions are annotated below. Alignment chains for MicOch1.0 and the meadow vole Hap1 assembly (this study) are shown at the bottom. (**D**) Dotplot of the annotated centromere. (**E**) Median CN of the two regions highlighted with red boxes; colors match the legend in (C). **(F)** Hap1 genomic alignments of the prairie vole SD comprising *Avpr1a* (purple) and *Avpr1a2* (red) with repeat annotations; the red box indicates the region enlarged at right. The hashed box upstream of both paralogs marks a microsatellite associated with altered *Avpr1a* expression^55,56^. (**G**) RNA-seq expression of *Avpr1a* paralogs in medial preoptic area (MPOA), dentate gyrus (DG), lateral septum (LS), nucleus accumbens (NAc), and subventricular zone (SVZ). Box plots show median (center line), interquartile range (box) and 1.5× IQR whiskers; dots are individual samples. TPM, transcripts per million reads. See also Figure S11–S16 and Tables S11–S18.

### Comparison of vole genomes identifies *Avpr1a* prairie-vole specific duplication

To better match orthologous regions, we re-oriented Hap1 for both vole assemblies based on synteny to the prairie vole reference (MicOch1.0) (Tables S6, S11, and S12). Alignment between the two species shows ∼98% similarity with notable cytogenetic differences, including several fusions and fissions, impacting 14 chromosomes in each genome (Figures 3A). Large-scale inversions are evident on several chromosomes, including homologous chromosomes 1 and X (Figure S13). Mapping single-nucleotide, indel, and structural variants identified ∼415K coding variants using prairie vole as a reference^34^; of these, 839 (VEP) and 1,043 (SnpEff) were annotated as likely gene disrupting (LGD) impacting a consensus set of 253 single-copy 1:1 ortholog genes (Tables S13 and S14). Focusing on candidate genes implicated in behavior—including those known to interact with nonapeptide hormones oxytocin and vasopressin—show global reduced rates of substitutions relative to synonymous mutations (Ka/Ks << 1; Figure S14) and no LGD variants obviously impacting function, suggesting purifying selection (Tables S15 and S16).

Understanding that gene duplication is a common mechanism of trait innovation across the animal kingdom^35^, we identified 306 and 389 gene duplications unique to prairie or meadow vole, respectively (Table S17). Intersecting with our candidate genes list, we identified a prairie-vole-specific duplication of *Avpr1a*, encoding vasopressin V1A receptor, a transmembrane G-protein coupled receptor that binds arginine vasopressin. While this duplication was characterized via BAC sequencing nearly 15 years ago^22^, both *Avpr1a* paralogs are missing in the current prairie vole reference (MicOch1.0) impeding analyses of these genes. Each prairie vole haplotype shows *Avpr1a* paralogs residing ∼900 kbp apart on chr26, separated proximally by a large stretch of nearly identical satellite repeats (547 kbp and 517 kbp) not present at the syntenic chr20 locus of the meadow vole (see chain alignment for Meadow vole Hap1; Figures 3B–3D). The *Avpr1a* paralogs flank the chr26 annotated metacentric centromere, defined as the largest stretch of satellite sequence per chromosome^36^ and including a >100-kbp hypomethylated region, not observed adjacent to *Avpr1a* in chr20 of the meadow vole (instead annotated as telocentric) nor at any other regions in either species’ genomes. Computing diploid copy number (CN) from Illumina read depth^37^ showed elevated CN across the chr26 centromere annotation and CN of 4 unique for the *Avpr1a*-containing SD (Figure 3E) in prairie vole versus CN of 2 in meadow vole, woodland vole (*M. pinetorum*, also known as pine vole, a species exhibiting monogamous pair bonding^38,39^ sequenced as part of this study), and an additional four publicly available *Microtus* genomes^40,41^ representing species with more promiscuous behaviors^42–47^.

Both curated prairie vole references show 97% nucleotide identity between *Avpr1a* paralogs, with *Avpr1a2* harboring frameshift variants encoding a truncated protein (218 aa vs. 420 aa full-length). This result was supported by both Hap1 and Hap2 assemblies (Figure S15 and Table S18) and is consistent with prior reports^21,22^. If translated, the first 199 aa would be identical between AVPR1A paralogs, encoding the extracellular vasopressin-binding domain and the first four of seven transmembrane domains but lacking the intracellular G-protein binding region^48^, with the terminating 19 aa unique to the prairie-vole-specific AVPR1A2. Quantifying transcript abundances using published RNA-seq data^49–54^ reveals consistent expression of both paralogs across five of eight tested brain regions, with the highest expression in the medial preoptic area, albeit reduced for *Avpr1a2* (∼1–4× relative to *Avpr1a*) (Figures 3G and S16). Divergent expression could be influenced by a 636-bp indel ∼1.6 kbp upstream of the *Avpr1a* start codons (Figure 3F) directly adjacent to a polymorphic microsatellite repeat previously associated with differences in *Avpr1a* expression and socio-behavioral traits in prairie voles^55,56^. Further, divergent chromatin interactions outside of the duplicated region likely occur between paralogs^57^ with the SD breakpoint only 4 kbp upstream. Even if *Avpr1a2* ultimately proves to be a non-functional pseudogene, the corrected assembly enables expression and epigenomic analysis of the ancestral full-length *Avpr1a* and its *cis*-regulatory landscape, previously absent from the prairie vole reference genome entirely.

## Discussion

Most recent efforts to build gigabase-sized chromosome-scale genomes require multiple technologies^58^, typically comprising highly accurate HiFi long reads (∼15-20 kbp at 30–60× coverage), ultralong nanopore reads (>100 kbp at ∼30× coverage), and short-read long-range information to phase and scaffold at chromosome scale^59^; even higher coverage is necessary to achieve complete diploid T2T genomes^60^. This “ideal” recipe is inaccessible to many researchers, requiring large amounts of starting material, multiple libraries, and three sequencing platforms. Here, we generated ∼100 Gbp of HiFi and CiFi data from a single library and sequencing run, producing contiguous (scaffold N50 90–125 Mbp) and high-quality diploid assemblies (QV>50). While large amounts of input were available here, experimental approaches are scalable to smaller amounts, as we previously demonstrated^7^. This is particularly important when studying rare, endangered, or difficult-to-sample species and tissues, where collecting large DNA quantities might be impractical. CiFi downsampling shows that 2× CiFi plus 30× HiFi coverage, achievable on a single SMRT Cell, is sufficient to accurately assemble a larger, more homogeneous (heterozygosity ∼0.1% vs 1% for voles) human diploid genome (3.2 Gbp) (Figures S5, S6 and Table S3). Relative to the “ideal” multi-platform recipe, the single-library approach reduces cost at least threefold while still resolving more than half of chromosomes at T2T scale. In effect, this lowers technical and cost barriers, enabling institutions with only a single sequencing platform to produce high-quality and contiguous genomes.

Beyond introducing a simplified assembly approach, we provide a long-awaited resource expanding the genomic toolkit for prairie voles, an emerging model organism for studying complex social behaviors relevant to humans^61^. This includes high-quality annotated assemblies for both prairie and meadow voles, repeat and SD annotations, CN maps across additional vole species, chain alignments facilitating genome liftover between species/builds (e.g., VGP mMicPen1), and named gene orthologs enabling transcriptomic comparisons, all publicly accessible through a UCSC Genome Browser hub (see STAR Methods). These resources complement ongoing neurobiological studies examining how social relationships are encoded in the brain.

Neuropeptide hormone pathways represent an important substrate through which neural circuits mediating innate behaviors can rapidly evolve and diversify^14,15^. Hence, divergence of the genes encoding neuropeptide receptors or their regulatory regions may underlie species differences in behavior. The highly contiguous assemblies generated here enabled discovery of a complex >350-kbp satellite repeat element with centromere annotation and confirmed the presence of a >100 kbp SD impacting *Avpr1a* (Figure 3). Neither feature is present in any of the four meadow vole haplotypes examined (this study and mMicPen1), nor was elevated CN detected at these regions in the five other vole species tested, including woodland vole, which also exhibits pair bonding. While this specificity to prairie vole argues against these variants as cross-species drivers of pair bonding, they remain compelling candidates for prairie-vole-specific functions. Hypo-methylation across satellite repeats strengthens annotation of the prairie vole chromosome 26 centromere and places *Avpr1a* approximately 350 kbp downstream, within the pericentromeric region. This positioning may expose the gene to centromerization-associated effects, including increased H3K27me3-mediated repression, altered chromatin accessibility, and elevated genomic instability^62^—consequences that would alter vasopressin signaling dynamics and likely facilitated the emergence of the prairie vole-specific *Avpr1a*2 paralog. This is a compelling finding given that prairie voles exhibit elevated *Avpr1a* expression in the ventral pallidum while meadow voles show higher expression in the lateral septum^21^. These assemblies will serve as an important resource for continued investigation of *Avpr1a* paralog functions and divergent brain expression patterns.

Our assessment of candidate behavioral genes identified no obvious amino-acid divergence between prairie and meadow voles (Figure S14 and Tables S15 and S16), suggesting that regulatory alterations, rather than coding sequence change, plays an outsized role in behavioral differences between these species. These near-complete genomes will enable comprehensive comparisons of altered chromatin structure and gene regulation underlying bond formation and social memory. Ultimately, integration with human genetic studies, such as *AVPR1A* variants linked to autism spectrum disorder^63–68^, provides a framework for understanding both how social behavior is encoded neurobiologically and how genetic disruptions to these pathways contribute to neuropsychiatric risk.

### Limitations of the study

A notable limitation is that current assembly and scaffolding tools, including hifiasm and YAHS, do not leverage the multi-contact information inherent to CiFi concatemers, instead requiring reduction into Hi-C-like pairs. Development of algorithms designed to leverage these higher-order chromatin contacts will likely yield further improvements in phasing, with CiFi showing an >8-fold greater ability to infer haplotypes versus Hi-C when considering multicontacts^7^, as well as scaffolding, particularly in complex genomic regions. Additionally, use of a single restriction enzyme (HindIII) introduces the possibility of sequence-biased coverage (also inherent in short-read Hi-C); adoption of alternative fragmentation approaches such as Omni-C could mitigate this in future implementations. Gene annotations might be further refined by incorporating long-read isoform sequencing, enabling a fully integrated workflow for assembly, scaffolding, and annotation from a single sequencing technology. Finally, although two haplotypes were resolved per species, single-individual sampling limits the ability to distinguish fixed from polymorphic differences. Broader sampling within and across *Microtus* species using a phylogenetic framework will be necessary to determine whether identified variants segregate with behavioral phenotypes. Such efforts would also support our observation of near-complete Y-chromosome divergence between prairie and meadow voles, in alignment with past work that suggests rapid chromosome evolution as a possible driver of the *Microtus* radiation^69^, one of the most dramatic speciation events recorded in vertebrates^70^.

In summary, we present a combined HiFi and CiFi sequencing strategy and simplified bioinformatic workflow enabling accurate, contiguous, chromosome-scale genome assembly from a single sequencing platform and experiment. Applying this approach to prairie and meadow voles, we generate high-quality diploid assemblies that reveal genome-wide differences between these behaviorally divergent species, with particular focus on loci implicated in social behavior. Ultimately, the simplicity, affordability, and minimal sample requirements of the combined HiFi–CiFi approach position it as a broadly accessible tool for comparative genomics, with the future potential to make chromosome-scale assembly tractable for rare, difficult-to-sample, and non-model organisms alike.

## Resource Availability

### Lead contact

Further information and requests for resources should be directed to and will be fulfilled by lead contact Megan Y. Dennis.

### Materials availability

● This study did not generate new unique reagents.

### Data and code availability

● All raw sequencing data generated from this study have been deposited to ENA and NCBI (PRJEB108798). Accession numbers of data used from published sources can also be found in the Key Resources Table.
● All source code, workflows, and processed data associated with this study is publicly available through https://github.com/mydennislab/2026-voles-assembly and https://github.com/sanger-tol/ and has been deposited at Zenodo (doi: 10.5281/zenodo.20501386^71^)
● Any additional information required to reanalyze the data reported in this paper is available from the lead contact upon request.

## Acknowledgments

We thank Dr. Aaron Wenger and Jacob Brandenburg for coordination support in generating and analyzing the HiFi-CiFi datasets. Also thanks to Drs. Michael Schatz and Daniela Soto for assistance with earlier iterations of vole genome assemblies, and Michael Sherman for help with animal husbandry. This work was supported, in part, by the U.S. National Science Foundation (CAREER 2145885 to M.Y.D.; CAREER IOS-2045348 to Z.R.D.) and the National Institutes of Health (NIH) grants from the National Institute of Mental Health (RF1MH132818 to M.Y.D. and DP2MH119427 to Z.R.D.). K.K., J.W., and K.H. are supported by Wellcome through the 220540/Z/20/A award that supports the Wellcome Sanger Institute. Voles and tissue samples were supported by NIMH (R01MH123513) and NSF (1556974) grants to D.S.M. CSHL voles studies were supported by an institutional grant to J.T. Woodland vole analysis was supported by a UC-Davis Interdisciplinary Catalyst Award to K.L.B. and M.Y.D. Illumina sequencing of RNA-seq and Hi-C libraries was performed at the Cold Spring Harbor Next Generation Sequencing Shared Resource, which is supported by NIH Cancer Center Support Grant 5P30CA045508 Some images were created using BioRender.

## Author Contributions

G.K., C.L., M.V.W., N.G.S., Z.R.D., K.L.B., and J.T. generated biospecimens and/or data for the project. M.A., J.L., K.K., J.W. K.H., and M.Y.D. performed bioinformatic and genomic analysis. K.L.B., D.M. and J.T. provided biomaterials and provided vole expertise. J.K., D.M., J.T., and M.Y.D. devised the project. M.A., J.K., D.M., and J.T., and M.Y.D. wrote the manuscript and all authors edited and approved the manuscript.

## Declaration of Interests

J.L. & C.L. are employees and shareholders and J.K. is a consultant and shareholder of Pacific Biosciences, a company developing single-molecule sequencing technologies. All other authors declare no competing interests.

## STAR Methods

### Experimental model and study participant details

#### Vole procedures

The male adult prairie vole (*M. ochrogaster*) and meadow vole (*M. pennsylvanicus*) individuals were maintained under protocols approved by the Institutional Animal Care and Use Committee (IACUC) at the University of California, San Francisco and Cold Spring Harbor Laboratories. Animals were euthanized according to institutional guidelines, and tissues were rapidly dissected, flash-frozen in liquid nitrogen, and stored at −80 °C until DNA or RNA extraction.

### Method details

#### HiFi and CiFi library preparations and sequencing of prairie and meadow vole liver samples

For whole-genome HiFi sequencing, high molecular weight (HMW) genomic DNA was extracted from frozen liver tissue using the Monarch HMW DNA Extraction Kit for Tissue (New England Biolabs) according to the manufacturer’s protocol. DNA quality and fragment size distribution were assessed by Qubit fluorometry and Femto Pulse analysis prior to downstream HiFi library preparation. For CiFi library preparation, frozen liver tissue (∼100 mg) was pulverized under liquid nitrogen and crosslinked, followed by processing according to CiFi protocol Part 1 (HindIII restriction digestion and proximity ligation)^7^. Crosslinks were reversed, and proximity-ligated 3C DNA was purified by phenol–chloroform extraction prior to downstream size selection and amplification.

A starting amount of 2.2 µg of HMW DNA was sheared on a Hamilton NGS STAR MOA system and size-selected on the Pippin HT with 10 kbp cutoff. The pre-PCR CiFi DNA (1.5 µg) was size-selected on the Pippin HT with 6.5 kbp cutoff and PCR amplified (50 ng input, 8 cycles) using the Ampli-Fi protocol (PacBio, 103-648-000). The post-PCR CiFi DNA was size-selected on the Pippin HT with 5.5 kbp cutoff. They were subsequently mixed at a molar ratio of 90% sheared DNA for HiFi and 10% CiFi DNA (translating to 650 ng and 35 ng, respectively, accounting for the size differences), followed by standard library preparation using SMRTbell prep kit 3.0 (PacBio, 102-182-700). The SMRTbell library was cleaned up with 1× SMRTbell cleanup beads. One SMRT Cell per species was run on the Revio system at 250 pM on-plate loading concentration with SPRQ chemistry and 30-hours acquisition.

HiFi and CiFi data from the sequencing run were segregated and adapters were trimmed with lima v2.14.0 (https://github.com/PacificBiosciences/barcoding) using a three-step process. First, the CiFi reads were identified by their unique dual index adapters and separated from HiFi reads using very relaxed lima settings (‘––ccs ––min-passes 0 ––min-end-score 0 ––min-score 5 ––min-ref-span 0.2 ––min-score-lead 0‘) to ensure that no CiFi reads remained in the HiFi data. Adapters were then trimmed from each dataset using the recommended settings: ‘––hifi-preset SYMMETRIC’ for the HiFi reads and ‘––hifi-preset ASYMMETRIC ––neighbors’ for the CiFi reads. PCR duplicates were then removed and duplication rates assessed for the CiFi reads using pbmarkdup v1.1.0 (https://github.com/PacificBiosciences/pbmarkdup).

#### Illumina sequencing vole genomic samples

Chromatin isolation and Hi-C library preparation was performed by Phase Genomics (WA, USA), using 100 mg flash-frozen liver tissue from the same individual male prairie and meadow voles as were used for HiFi and CiFi library preparation. The Proximo Hi-C protocol (Phase Genomics, WA, USA)^73^ was used to prepare the proximity ligation library and processed into an Illumina-compatible sequencing library. Hi-C libraries were sequenced with 150-bp paired-end reads on an Illumina NextSeq500 at the CSHL Next-Generation Sequencing Core Facility.

gDNA was extracted from dried museum pelt tissue of the woodland vole (*M. pinetorum*) using the DNeasy Blood & Tissue Kit (Qiagen) following manufacturer instructions with minor modifications to reduce PCR inhibitors. Extracted DNA was treated with RNase A and further purified by ethanol precipitation. Sequencing libraries were prepared for short-read sequencing using prepared gDNA for prairie vole, meadow vole, and woodland vole. Prepared libraries were sequenced (2×150 bp) by Novogene on a NovaSeq 6000 platform to produce approximately 60 Gbp of raw data (∼30× genome coverage) for each species.

#### Genome assembly and scaffolding

HiFi reads were extracted from unaligned BAM files using samtools v1.21^74^ (‘samtools fastà). CiFi reads were converted from BAM to FASTQ (‘samtools collate –O –u | samtools fastq’) and then processed into Hi-C-like paired-end reads using the cifi-toolkit (https://github.com/mydennislab/cifi-toolkit) that performs *in silico* HindIII digestion on each CiFi concatemer read, extracting the outermost restriction fragments as R1 and R2 paired-end reads. Phased diploid assembly was performed with hifiasm v0.19.8^4^ using the ‘––dual-scaf’ mode, which integrated either CiFi (input as paired-end reads) or Hi-C contact information during graph resolution to produce haplotype-resolved primary contig graphs for each haplotype (Hap1 and Hap2). HiFi FASTA and derived R1/R2 FASTQ files were provided as input (‘––h1’, ‘––h2’). GFA contig graphs were converted to FASTA using gfatools v0.5 (https://github.com/lh3/gfatools).

CiFi and Hi-C contact maps were generated by aligning the full CiFi BAM to each haplotype assembly using the epi2me-labs/wf-pore-c Nextflow pipeline v1.3.0 (https://github.com/epi2me-labs/wf-pore-c) with minimap2 in ‘-ax map-hifì mode, HindIII as the restriction enzyme, and ‘––paired_end’ enabled to produce BED contact files. Contig scaffolding was performed with YAHS v1.2a.2^23^ using the Pore-C BED contact file as input, with contig error correction disabled (‘––no-contig-ec’).

To evaluate the minimum CiFi sequencing depth required for chromosome-scale scaffolding, the CiFi BAM was subsampled at 1%, 10%, 15%, 20%, 25%, 40%, 60%, 80%, and 100% of the original read depth using ‘samtools view –s’ with a fixed random seed. At each fraction, the full assembly and scaffolding pipeline described above was executed independently, and scaffold N50 and auN were compared across titration points.

#### NA12878 CiFi haplotype phasing benchmark

Because no phased truth set exists for the vole species sequenced in this study, phasing accuracy was benchmarked on the human GIAB HG001/NA12878 sample. Public PacBio HiFi reads from NA12878^24^ were paired with CiFi reads (prepared using HindIII) from the matched GM12878 lymphoblastoid line^7^. HiFi reads were downsampled to 30× on GRCh38, and CiFi reads were downsampled to 0, 2, 3.5, 5, 7.5, 10, and 15× and also evaluated at full depth. A matched Hi-C analysis used a public GM12878 Hi-C dataset (prepared with MboI)^75^ at the same coverage (2 to 30×). CiFi reads were converted to paired-end concatemer segments by in silico digestion (‘cifi digest’). For each condition, hifiasm v0.25.0 was run on the 30× HiFi reads with paired-end CiFi or Hi-C reads supplied via ‘––h1’/‘––h2’ (‘––dual-scaf ––telo-m CCCTAAÀ); the 0×-CiFi baseline used hifiasm on HiFi alone. Each phased haplotype was aligned to GRCh38 with dipcall v0.3, and ‘whatshap comparè (v2.7) was run on the dipcall × GIAB confident intersection (HG001 v4.2.1, autosomes chr1–chr22) to compute per-chromosome switch error and blockwise Hamming error rates and against the pedigree-phased NA12878 call set (Platinum Pedigree^24^).

#### Genome assembly manual curation

To facilitate manual assembly curation for each genome of the prairie and the meadow voles, the two scaffolded haplotypes were combined together and the corresponding CiFi datasets were mapped onto the assemblies using wf-pore-c pipeline with parameters ‘––paired_end ––cutter HindIII ––paired_end_minimum_distance 100 ––paired_end_maximum_distance 200’. Chromatin contact maps were further generated for the combined haplotypes of each genome with PretextMap using the mock paired-end BAM files as input and retaining non-uniquely mapping reads in the contact maps with –mapq 0. Supplementary analysis were also embedded in the Pretext file using the Tree of Life curationpretext pipeline – telomeres with standard vertebrate motif TTAGGG, N base scaffold gaps and mapped PacBio long-read coverage. AGP files of the corrected contact maps were exported from PretextView and curated haplotype assemblies generated using pretext-to-asm.

#### Completeness and contiguity assessment

To evaluate the contiguity and completeness of our assemblies, we assessed telomere presence and sequence gaps across all chromosome-scale scaffolds for both haplotypes of each species. Gaps (runs of N bases) were identified using seqtk. Telomeric repeat sequences were detected using tidk v0.2 (Telomere Identification Toolkit;^76^), which scans for the canonical vertebrate telomere motif (TTAGGG) in sliding windows across each scaffold. We searched for TTAGGG repeats in 10-kbp windows and considered a telomere present at a chromosome terminus when the terminal window contained at least 10 repeat units (forward and reverse strands combined). Chromosomes were classified as T2T (telomeric repeats at both ends), partial (one end only), or incomplete (neither end). A chromosome was considered T2T scaffolds when both telomere ends were detected but with sequence gaps were present. A fully T2T assembly comprised both telomere ends and a single contig with no gaps. Genome completeness of each assembly was also assessed with BUSCO v6.0.0^77^ in genome mode against the glires_odb12 lineage (12,556 orthologs)^78^.

#### Heterozygosity and consensus quality estimation

Per-individual heterozygosity and consensus quality (QV) were estimated from Illumina paired-end whole-genome sequencing of gDNA prepared from the same individuals used for HiFi sequencing. Per-species k-mer databases were built with Meryl v1.3^25^ using k=21. Heterozygosity was estimated by extracting the k-mer frequency histograms from the Meryl databases (meryl histogram) and fitting a diploid mixture model with GenomeScope2 v2.1.0^79^ and k-mer completeness were computed with Merqury v1.3 using k=31, with diploid values represented in Table 1. Haplotype-specific analyses provided similar QV (prairie vole 51.9 (Hap1) and 52.0 (Hap2); meadow vole 50.0 (Hap 1) and 52.1 (Hap2)) and k-mer completeness estimates (prairie vole 83.3% (Hap1) and 78.6% (Hap2); meadow vole 81.2% (Hap 1) and 76.6% (Hap2)).

**Table 1.**
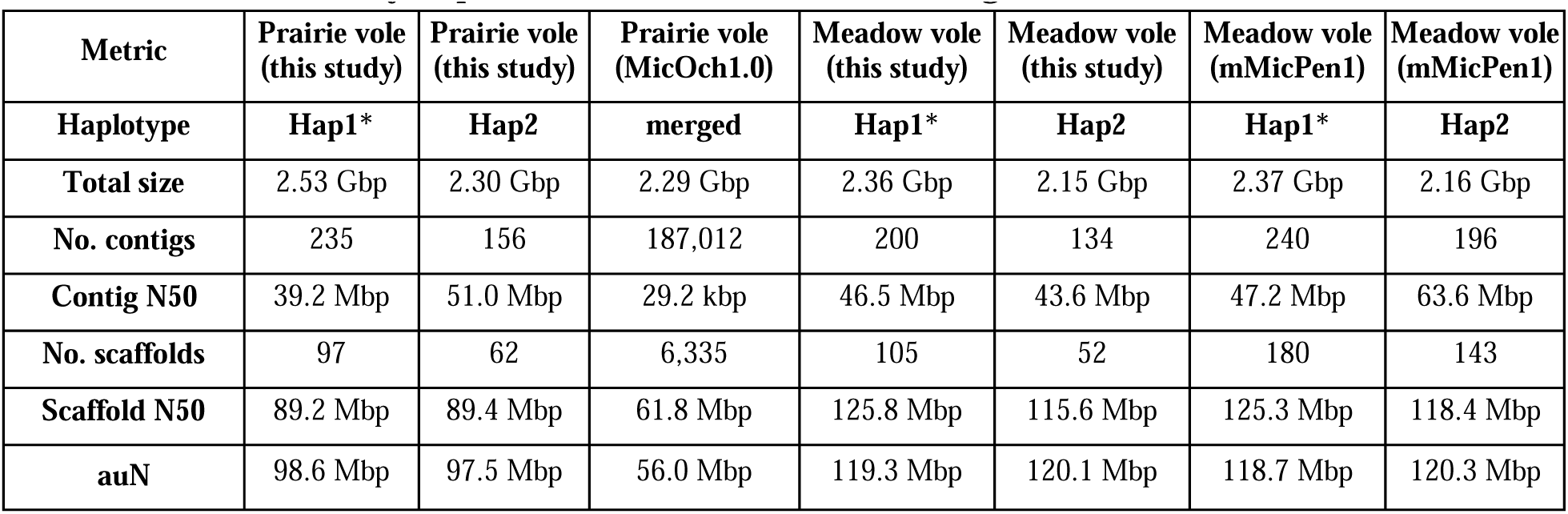
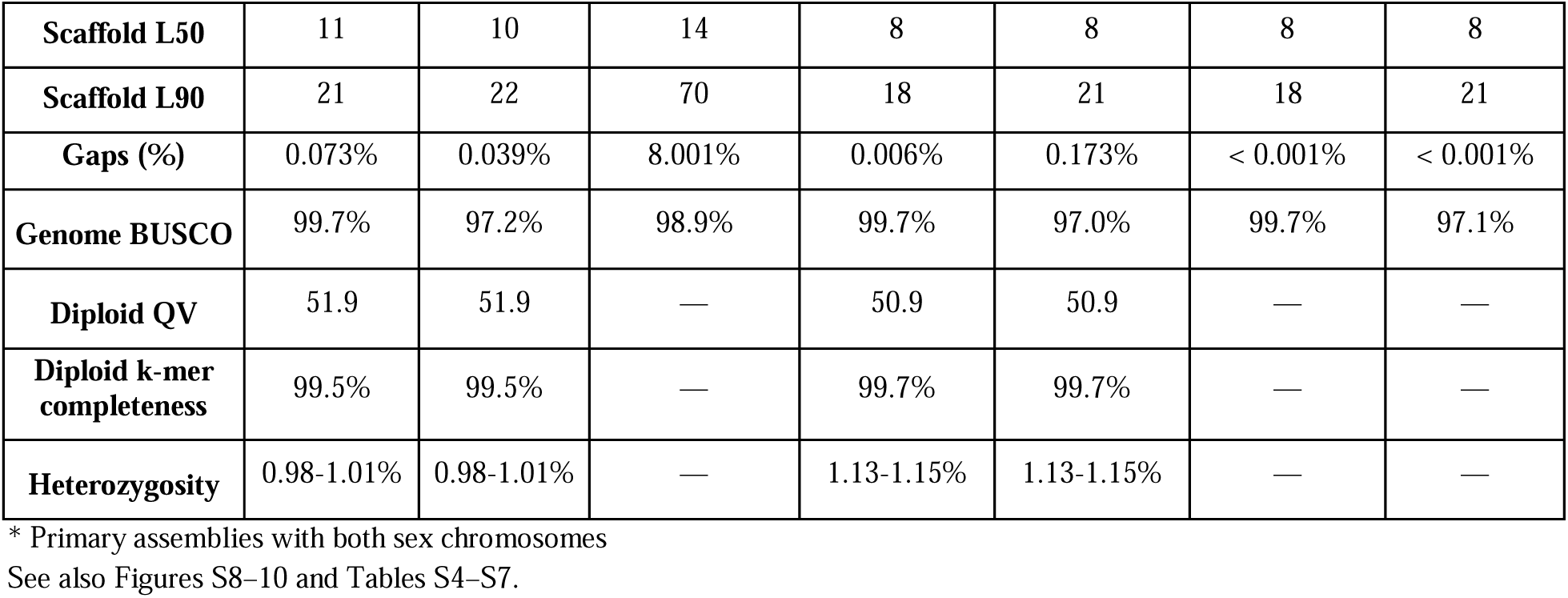
Metric summary of prairie and meadow vole curated genomes.

| Metric | Prairie vole (this study) | Prairie vole (this study) | Prairie vole (MicOch1.0) | Meadow vole (this study) | Meadow vole (this study) | Meadow vole (mMicPen1) | Meadow vole (mMicPen1) |
| --- | --- | --- | --- | --- | --- | --- | --- |
| Haplotype | Hap1* | Hap2 | merged | Hap1* | Hap2 | Hap1* | Hap2 |
| Total size | 2.53 Gbp | 2.30 Gbp | 2.29 Gbp | 2.36 Gbp | 2.15 Gbp | 2.37 Gbp | 2.16 Gbp |
| No. contigs | 235 | 156 | 187,012 | 200 | 134 | 240 | 196 |
| Contig N50 | 39.2 Mbp | 51.0 Mbp | 29.2 kbp | 46.5 Mbp | 43.6 Mbp | 47.2 Mbp | 63.6 Mbp |
| No. scaffolds | 97 | 62 | 6,335 | 105 | 52 | 180 | 143 |
| Scaffold N50 | 89.2 Mbp | 89.4 Mbp | 61.8 Mbp | 125.8 Mbp | 115.6 Mbp | 125.3 Mbp | 118.4 Mbp |
| auN | 98.6 Mbp | 97.5 Mbp | 56.0 Mbp | 119.3 Mbp | 120.1 Mbp | 118.7 Mbp | 120.3 Mbp |
| **Scaffold L50** | 11 | 10 | 14 | 8 | 8 | 8 | 8 |
| **Scaffold L90** | 21 | 22 | 70 | 18 | 21 | 18 | 21 |
| **Gaps (%)** | 0.073% | 0.039% | 8.001% | 0.006% | 0.173% | < 0.001% | < 0.001% |
| **Genome BUSCO** | 99.7% | 97.2% | 98.9% | 99.7% | 97.0% | 99.7% | 97.1% |
| **Diploid QV** | 51.9 | 51.9 | — | 50.9 | 50.9 | — | — |
| **Diploid k-mer completeness** | 99.5% | 99.5% | — | 99.7% | 99.7% | — | — |
| **Heterozygosity** | 0.98-1.01% | 0.98-1.01% | — | 1.13-1.15% | 1.13-1.15% | — | — |

\* Primary assemblies with both sex chromosomes See also Figures S8–10 and Tables S4–S7.

#### Centromere annotation

Centromeric regions were identified on each Hap1 primary assembly using centroAnno v1.0.2^36^ in assembly annotation mode (‘-x anno-asm’). Each chromosome was processed individually; centroAnno decomposed tandem satellite repeats into monomer units and reported their genomic coordinates. Contiguous monomer annotations separated by fewer than 10 kbp were merged into repeat regions, and the largest repeat region per chromosome was designated as the putative centromere. Centromeres were detected on all chromosomes of both species. The sequence comprising the chr26 centromere using minimap (default parameters) matched satellite repeat sequences on chromosomes 4 and X (∼500 bp to 3 kbp in size at <93% sequence identity) for both Hap1 and Hap2 meadow vole assemblies but is found only at the chromosome 26 *Avpr1a* locus in both prairie vole haplotypes.

#### HiFi read mapping and methylation analysis

PacBio Revio HiFi reads carrying per-base 5mC calls (MM/ML BAM tags from the Jasmine primary-analysis basecaller) were processed with nf-core/methylong v2.0.0^80^ against the prairie and meadow vole references. Per-CpG bedMethyl outputs were converted to per-site and 1-kbp-windowed bigWig tracks with UCSC bedGraphToBigWig. Region-scale methylation segmentation into high– and low-methylated blocks was performed with MethBat as part of the nf-core/methylong workflow and rendered as a bigBed track; these segments classify each block as methylated or unmethylated within a sample. Coverage tracks were generated with mosdepth^81^ (1-kbp windows) on the HiFi methylation BAMs and on CiFi alignments produced by minimap2 v2.30 (map-hifi preset).

#### Segmental duplication and repeat annotations

Species-specific *de novo* repeat libraries were constructed for each vole species using RepeatModeler v2.0.7^82^ with default parameters. A RepeatModeler database was built from each Hap1 primary assembly, and *de novo* repeat family consensus sequences were identified. The resulting species-specific libraries were combined with the Dfam database and used as input for RepeatMasker v4.2.2^83^ with the RMBLAST search engine. RepeatMasker was executed via the Dfam TE Tools docker container; RepeatModeler v2.0.7, RepeatMasker v4.2.1, RMBLAST v2.14.1+). SDs were identified using BISER v1.4^31^ with default parameters. BISER was run on the soft-masked Hap1 assemblies for each species, and output was generated in BEDPE format.

Dot plots were generated using ODP v0.3.3^72^, circular genome visualizations using pyCirclize v1.9.1, and linear karyotype ideograms using matplotlib v3.10.8.

#### RNA-seq of vole tissues

Total RNA was prepared from flash frozen prairie and meadow vole tissues: liver, brain, and testes from adults and e11.5 (prairie) or e13.5 (meadow) embryos. 50–75 mg of tissue per sample was homogenized in 500 µl Trizol (Life Technologies) on ice using a Kimble Kontes Disposable Pellet Pestle (VWR). An additional 500 μl Trizol was added followed by further homogenization using an 18-gauge needle on a 1-ml syringe before proceeding to RNA extraction according to the Trizol protocol. The subsequent RNA was DNase treated with a TURBO DNA-free kit (Life Technologies). 150 ng of total RNA was used as input for library preparation with Encore Complete RNA-seq kits (NuGen), using ten cycles of amplification. Multiplexed libraries were sequenced with 76-bp paired-end reads on the Illumina NextSeq 500 at the CSHL Next-Generation Sequencing Core Facility.

#### Gene annotation

Illumina RNA-seq data from the four samples per species were used in the NCBI Eukaryotic Genome Annotation Pipeline (EGAPx) v0.4.1-alpha^84^ to perform gene annotation for both Hap1 assemblies of both species.

#### Gene expression analysis

Transcript quantification was performed using Salmon v1.10.3^85^ in mapping-based mode. For each species, *Avpr1a* transcript-level expression was first quantified across four in-house paired-end Illumina RNA-seq libraries (brain, liver, testes, and embryo) using a targeted Salmon index containing the *Avpr1a* coding sequences with the Hap1 genome assembly as a decoy. For the prairie vole, the index included both *Avpr1a* paralogs to enable paralog-specific quantification. To contextualize *Avpr1a* expression within the broader transcriptome, we performed transcriptome-wide Salmon quantification for the prairie vole. The full EGAPx-annotated transcriptome was extracted from the prairie Hap1 assembly using gffread, and a Salmon index was built by concatenating the transcriptome with the genome assembly. We quantified 453 publicly available prairie vole brain RNA-seq libraries from previously published studies^49–54^ spanning eight brain regions—amygdala (AMY), dentate gyrus (DG), hypothalamus (HT), lateral septum (LS), medial preoptic area (MPOA), nucleus accumbens (NAc), subventricular zone (SVZ), and ventral pallidum (VP)—under BioProject accessions PRJNA428754, PRJNA682808, PRJNA631040, PRJNA786347, PRJNA887096, PRJNA792575, PRJNA1005323, and PRJEB89367. After excluding 147 technical replicates, 306 samples were retained for analysis. Salmon was run with ‘––validateMappings’, ‘––gcBias’, and ‘––seqBias’ correction, with library type automatically inferred.

#### Scaffold orientation

To standardize chromosome naming, chromosome-scale scaffolds in the prairie Hap1 assemblies were renamed from SUPER_1–SUPER_26, SUPER_X, and SUPER_Y to chr1–chr26, chrX, and chrY, respectively. Scaffold numbering was retained from the *de novo* assembly. Scaffold orientation was then evaluated relative to the MicOch1.0 prairie vole reference assembly (GCF_000317375.1). Whole-genome alignments were generated with minimap2 v2.30 using –cx asm5 ––cs in both query-to-reference and reference-to-query directions, and alignment blocks shorter than 10 kbp were excluded. For each scaffold, orientation was inferred from the dominant aligned strand based on aligned bases; scaffolds with >50% of aligned bases on the reverse strand were reverse-complemented. Orientation confidence was classified as high when ≥95% of aligned bases supported the dominant strand, medium when 80–95% supported the dominant strand, and low/mixed when <80% supported the dominant strand. Because MicOch1.0 was generated from a female individual and lacks chrY, the prairie chrY scaffold was retained in its native orientation.

Prairie scaffold orientations were cross-validated using the published prairie vole genetic linkage map^27^. Marker flanking sequences were aligned to the assembly with BLASTn using an e-value threshold of 1 × 10^-10^, retaining the single best hit per marker. For each linkage group, the Spearman rank correlation between genetic position in centimorgans and physical position in the assembly was calculated. Linkage-map evidence was used to override the alignment-based orientation only when the alignment call had low/mixed confidence, |ρ| ≥ 0.70, and ≥5 markers were mapped to the scaffold. Discordant cases are reported in Table S6.

The corrected prairie Hap1 assembly was generated by reverse-complementing scaffolds designated by this procedure. Corrected scaffolds were then re-aligned to MicOch1.0 to record post-correction forward-strand percentages. Scaffolds with persistently low forward-strand alignment after correction were flagged in Tables S6 and S11, consistent with mosaic content or internal inversions that cannot be resolved by a single scaffold-level reverse-complement. A UCSC liftOver chain was generated to convert coordinates between the original and orientation-corrected assemblies.

To facilitate cross-species comparative analysis, meadow vole Hap1 scaffolds were oriented against the orientation-corrected prairie vole Hap1 assembly using the same alignment-based procedure, with minimap2 run using the cross-species preset –cx asm10. Meadow chrY had no reliable cross-species alignment and was retained in its native orientation. Minor meadow scaffolds were oriented when sufficient cross-species alignment evidence was available; scaffolds without informative alignments were retained in their native orientation. Per-scaffold orientation calls, alignment statistics, linkage-map validation results, and coordinate-conversion resources are provided in Tables S6, S7, S11, and S12.

To enable coordinate conversion from our meadow vole Hap1 assembly to the published VGP meadow vole reference (mMicPen1.hap1; GCF_037038515.1), we generated a UCSC liftOver chain file by aligning the two meadow vole assemblies. Our Hap1 assembly scaffolds were aligned to mMicPen1.hap1 using minimap2 v2.30 (‘-cx asm5 ––cs’). The resulting alignment was converted to UCSC chain format with transanno v0.4.5 ‘minimap2chain’^86^ and sorted with UCSC ‘chainSort’^87^. The liftOver chain file is provided as Supplementary Data and is compatible with UCSC ‘liftOver’ and related coordinate-conversion tools.

#### AVPR1A amino-acid alignment

Coding sequences for the prairie and meadow vole *Avpr1a* homologs were compared across both haplotypes: six sequences in total, comprising prairie *Avpr1a* and *Avpr1a2* and meadow *Avpr1a*, each from Hap1 and Hap2. Coding sequences were taken from the EGAPx gene annotations^88^ of each assembly; because the Hap2 assemblies were unannotated, the hap1 *Avpr1a* proteins were aligned to the Hap2 contigs with miniprot^89^ and the coding sequences extracted from the resulting gene models. The six coding sequences were aligned with MACSE v2^90^, a codon-aware aligner that accommodates the frameshift and internal stop codons of the *Avpr1a2* paralog and produces a corresponding amino-acid alignment.

#### Comparative assembly and paralog analyses

One-to-one orthologs between meadow and prairie voles were identified with OrthoFinder v2.5.5^91^ from the species’ EGAPx proteomes. The meadow vole Hap1 assembly was aligned to the prairie vole Hap1 reference with minimap2 v2.30^92^ (asm10 preset), and variants were called with paftools.js (alignment blocks ≥10 kb, MAPQ ≥30); calls were restricted to autosomal coding sequence and normalized with bcftools v1.22^74^. Coding variants were annotated independently with SnpEff v5.4c^93^ and Ensembl VEP v114.1^34^, and were retained as candidate gene-disrupting if any consequence carried IMPACT = HIGH and matched the high-impact Sequence Ontology set (stop_gained, frameshift_variant, splice_acceptor_variant, splice_donor_variant, start_lost, stop_lost, transcript_ablation, exon_loss_variant). Indels within 1 bp of a reference mononucleotide run of ≥6 bp were flagged as homopolymer artefacts and excluded from the per-gene evidence count; candidates were further restricted to 1:1 ortholog groups with a canonical gene symbol, each retained group required at least two distinct non-homopolymer HIGH-impact variants, and genes called by both annotators defined the final candidate-LoF set. Pairwise meadow-prairie Ka/Ks was computed in BioPython v1.86^94^ for every 1:1 ortholog carrying at least one coding SNV by applying the VEP-annotated substitutions to the prairie reference coding sequence and scoring each gene by the Nei-Gojobori (1986) method^95^. Pairwise comparisons and visualization of homologous chromosomes, as well as SD sequence containing *Avpr1a* (prairie vole Hap1 chr26:13,970,803-14,087,830) and *Avpr1a2* (prairie vole Hap1 chr26:13,058,418-13,176,477) were performed using MUMmer4 alignments^96^ and visualized with pyGenomeViz v1.6.1^97^. The annotated centromeric sequence (prairie vole Hap1 chr26:13,182,318-13,728,901) was queried against both Hap1 and Hap2 of the prairie and meadow vole assemblies (this study and mMicPen1) using minimap2 (v2.26)^92^ selecting for matches 90% or higher. Dot plots for chromosomes Y (both prairie and meadow voles) and 23 (prairie vole) were generated using nf-core/pairgenomealign^98^.

#### Copy-number analysis

Copy number (CN) was estimated using the FastCN pipeline^37^ with mrsFAST v3.4.2. All species were mapped to the prairie vole Hap1 primary assembly as a common reference to enable direct cross-species comparison at the same genomic coordinates. A four-layer masked reference was constructed from the prairie vole Hap1 assembly. Repetitive elements were masked using the RepeatMasker annotations described above, tandem repeats were identified with Tandem Repeats Finder^99^ per chromosome, and low-complexity regions were masked with WindowMasker/DUST^100^. The masked genome was then indexed with mrsFAST and all 50-mers were extracted and searched back against the genome; positions where a 50-mer aligned 20 or more times were additionally masked (K50 masking). Gaps in the final masked reference were extended by 36 bp on each side to account for the read-length shadow effect. Control regions expected to be diploid (CN=2) were defined by excluding SDs (identified by BISER), centromeric regions, and target gene loci from the genome-wide window set.

For each species, the first 36 bp were extracted from each mate of the paired-end whole-genome sequencing reads and mapped to the masked reference using mrsFAST with up to two mismatches allowed. GC-corrected read depth was computed in 1-kbp windows across the genome. CN was calculated as CN = 2 x (window depth / autosomal control mean). To prevent inflated CN values from masked regions with zero depth falling within control windows, we excluded zero-depth windows from the control mean calculation. Control regions were further refined by retaining only windows where all samples showed consistent CN near 2.0 (coefficient of variation < 0.2), and a final normalization ensured the median CN at refined control windows equaled 2.0.

Whole-genome sequencing reads from seven *Microtus* species—prairie vole (*M. ochrogaster*, this study), meadow vole (*M. pennsylvanicus*, this study), woodland vole (pine vole, *M. pinetorum*, this study), montane vole (*M. montanus*; SRR12966109), common vole (*M. arvalis*; ERR3427942), field vole (*M. agrestis*; SRR2167807), North American water vole (*M. richardsoni*; SRR12963053)— were mapped to the prairie vole reference. Each species was then normalized independently by scaling CN values so the median at refined diploid autosomal control regions equals 2.0, correcting for sequencing depth differences between species. Per-gene CN was calculated as the mean CN across all non-zero 1-kbp windows overlapping each target locus. CN evaluated for the putative centromere (prairie vole Hap 1, chr26:13,182,318-13,728,901), where estimates tend to be unreliable due to high repetitive content producing repeat masking, showed elevated CN in prairie vole across the chr26 centromere annotation relative to all other voles, suggesting this region is prairie-vole specific. CN estimation of the *Avpr1a* 118-kbp SD region (prairie vole Hap 1, chr26:13,970,803-14,087,830) confirmed the duplication to be prairie-vole specific with diploid CN of 4 compared to CN of 2 for all other tested vole genomes.

### Quantification and statistical analysis

Quantification procedures are described in the corresponding Method details sections. Analyses were primarily descriptive and are reported as counts, proportions, medians, or genome-wide/interval-level summary metrics unless otherwise stated. No additional formal statistical hypothesis testing was performed beyond the quantitative metrics described in the STAR Methods. For RNA-seq box plots, center lines indicate medians, boxes indicate interquartile ranges, whiskers indicate 1.5× the interquartile range, and points indicate individual samples.

### Additional resources

● UCSC Genome Browser hub for prairie vole: https://genome.ucsc.edu/cgi-bin/hgTracks?hgS_doOtherUser=submit&hgS_otherUserName=mabuelanin&hgS_otherUserSessionName=prairie_vole_cifi_hifi_hap1
● UCSC Genome Browser hub for meadow vole: https://genome.ucsc.edu/cgi-bin/hgTracks?hgS_doOtherUser=submit&hgS_otherUserName=mabuelanin&hgS_otherUserSessionName=meadow_vole_cifi_hifi_hap1
● General information and resources related to this project can be found at dennislab.org/voles

## Supplemental Information Index

Document S1. Figures S1–S16, Tables S1, S2, S4, S6, S8, S9, S10, and S13, Supplemental methods, and supplemental references.

Table S3. GM12878 haplotype phasing benchmark, related to Figure 1.

Table S5. Telomere-to-telomere (T2T) status of vole assemblies, related to Table 1.

Table S7. Prairie vole assembly resolved unplaced scaffolds, related to Table 1.

Table S11. Prairie scaffolds orientation decisions, related to Figure 3.

Table S12. Meadow scaffolds orientation decisions, related to Figure 3.

Table S14. Meadow vole loss-of-function candidate genes, related to Figure 3.

Table S15. Nucleotide divergence of behavior-related genes between voles, related to Figure 3.

Table S16. Variants distinguishing meadow and prairie voles across candidate behavior genes, related to Figure 3.

Table S17. Species-specific gene duplications, related to Figure 3.

Table S18. Avpr1a paralogs comparative analysis across species and haplotypes, related to Figure 3.

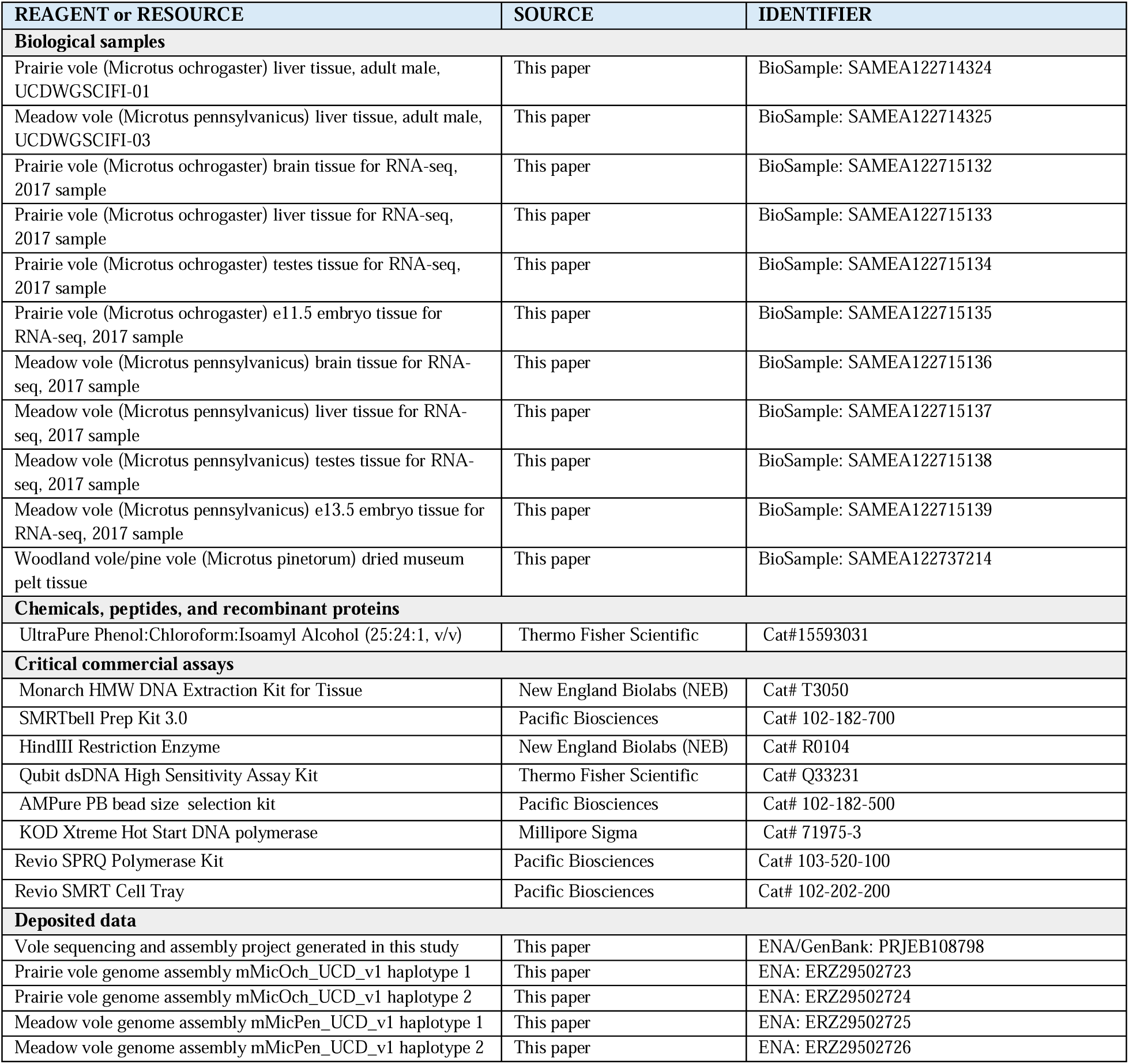
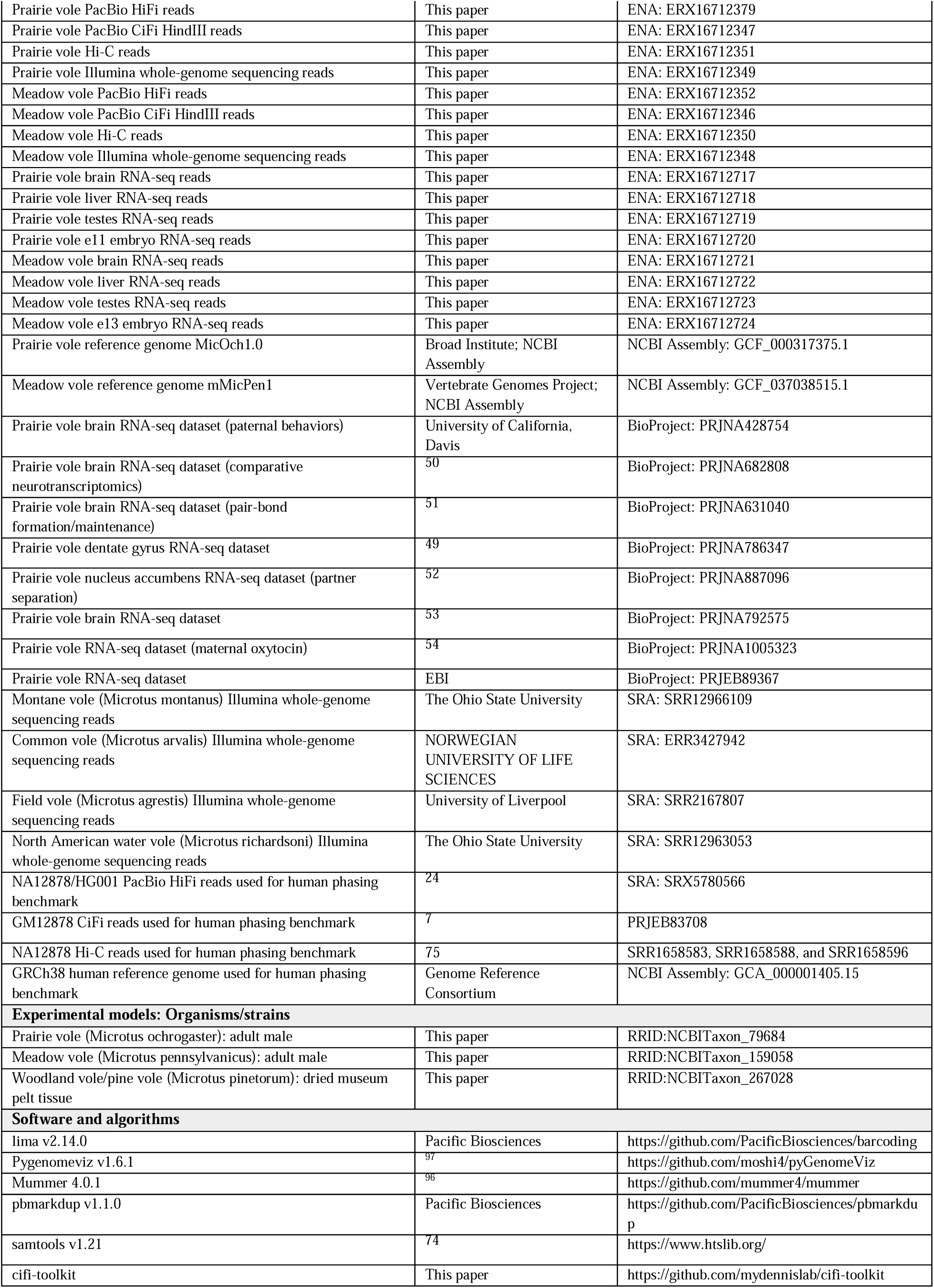
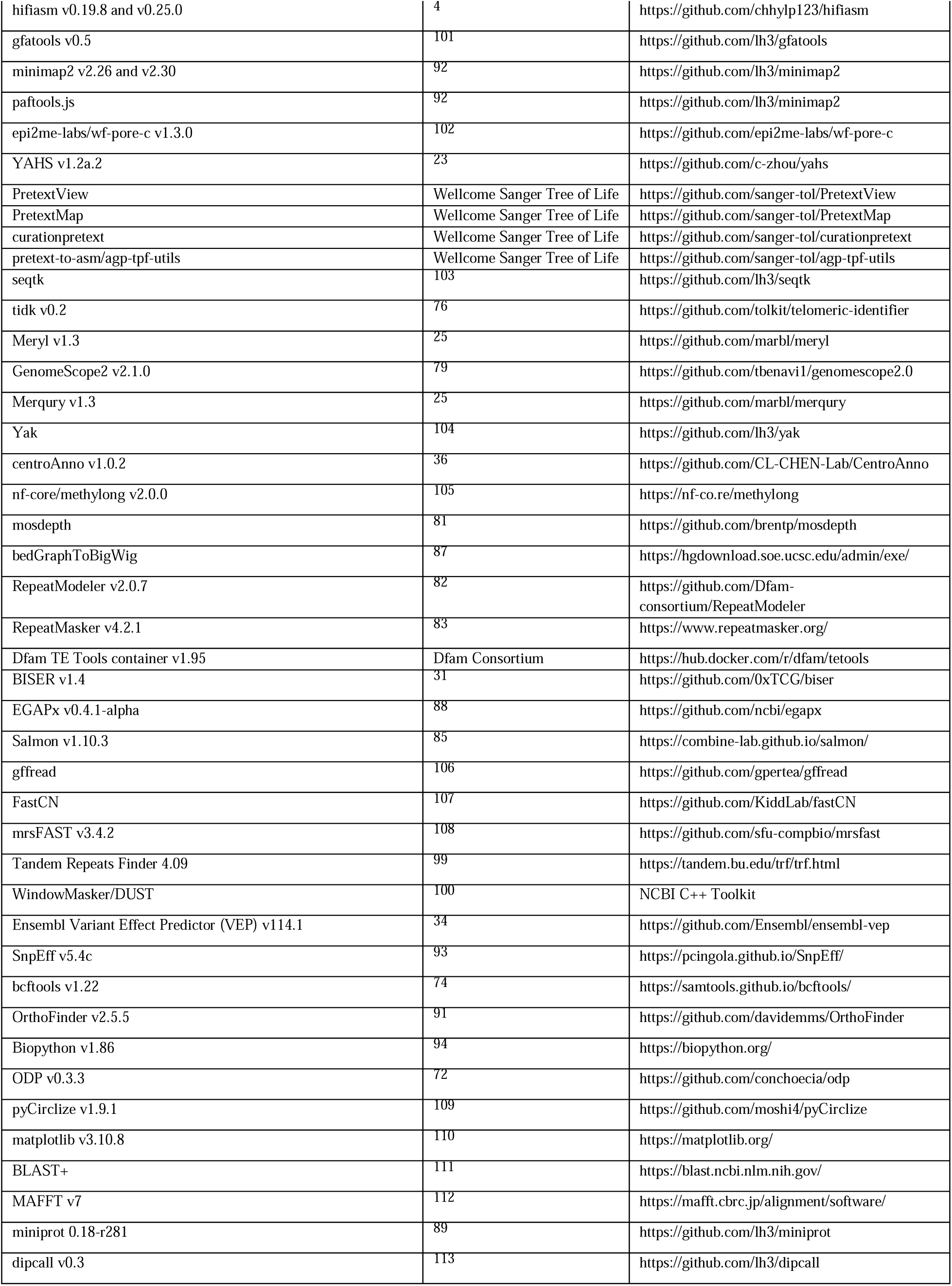
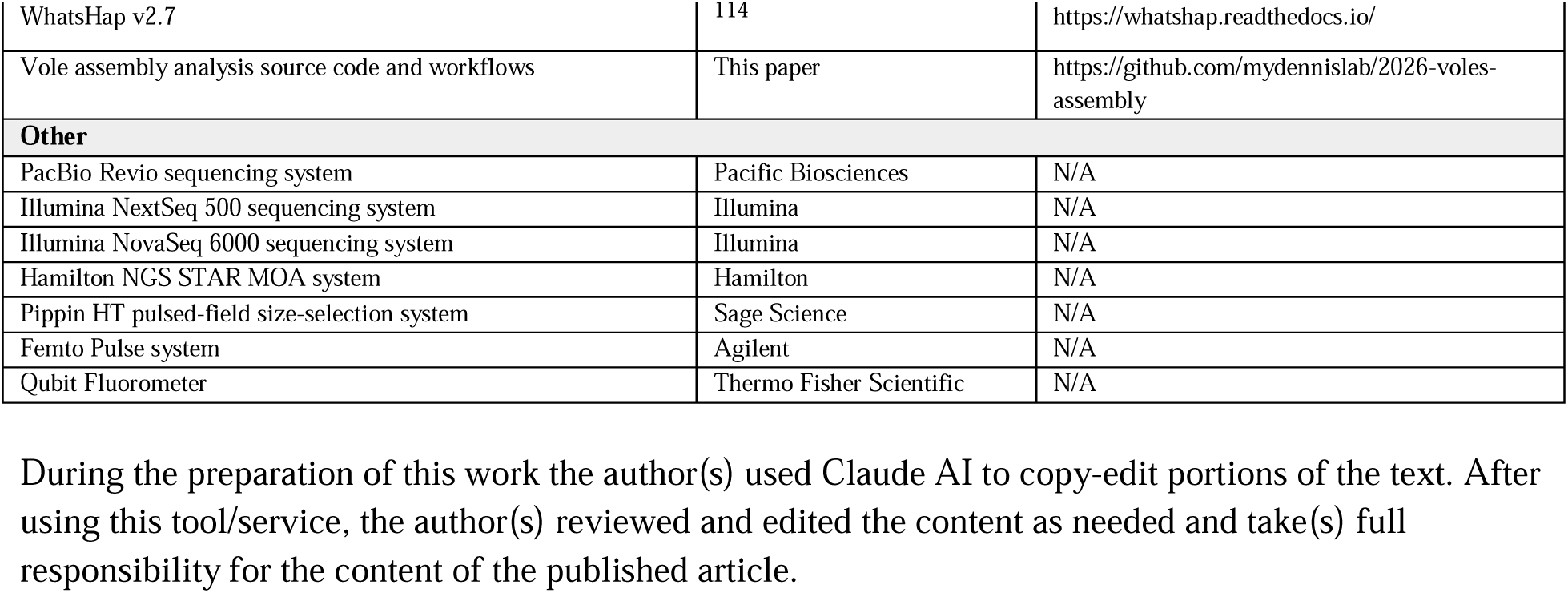
Key Resources Table.

## Notes

https://github.com/mydennislab/2026-voles-assembly

https://dennislab.org/voles

### Summary of Updates

Abstract simplified and shortened, added new analyses including benchmarking haplotype-phasing accuracy for human NA12878 dataset, methylation assessment, and updated species comparisons of disruptive variants.

