## Supplementary material for "Single-library chromosome-scale diploid assemblies of vole genomes resolve a species-specific duplication implicated in pair bonding": Document S1

### Supplemental Figures

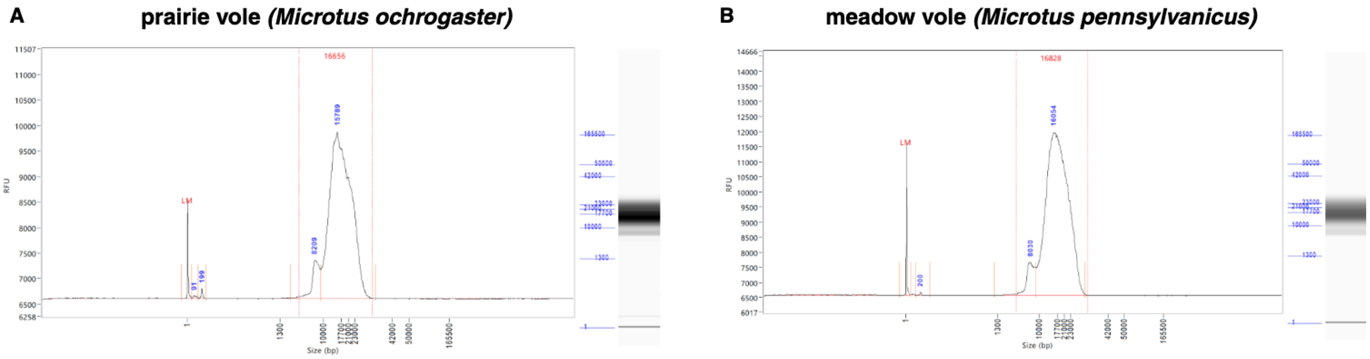

**Figure S1. Final pooled SMRTbell library size distributions, related to Figure 1.** Femto Pulse was used to assess final pooled HiFi and CiFi SMRTbell libraries for **(A)** prairie vole and **(B)** meadow vole. Size distribution profiles confirm expected fragment sizes prior to PacBio Revio sequencing.

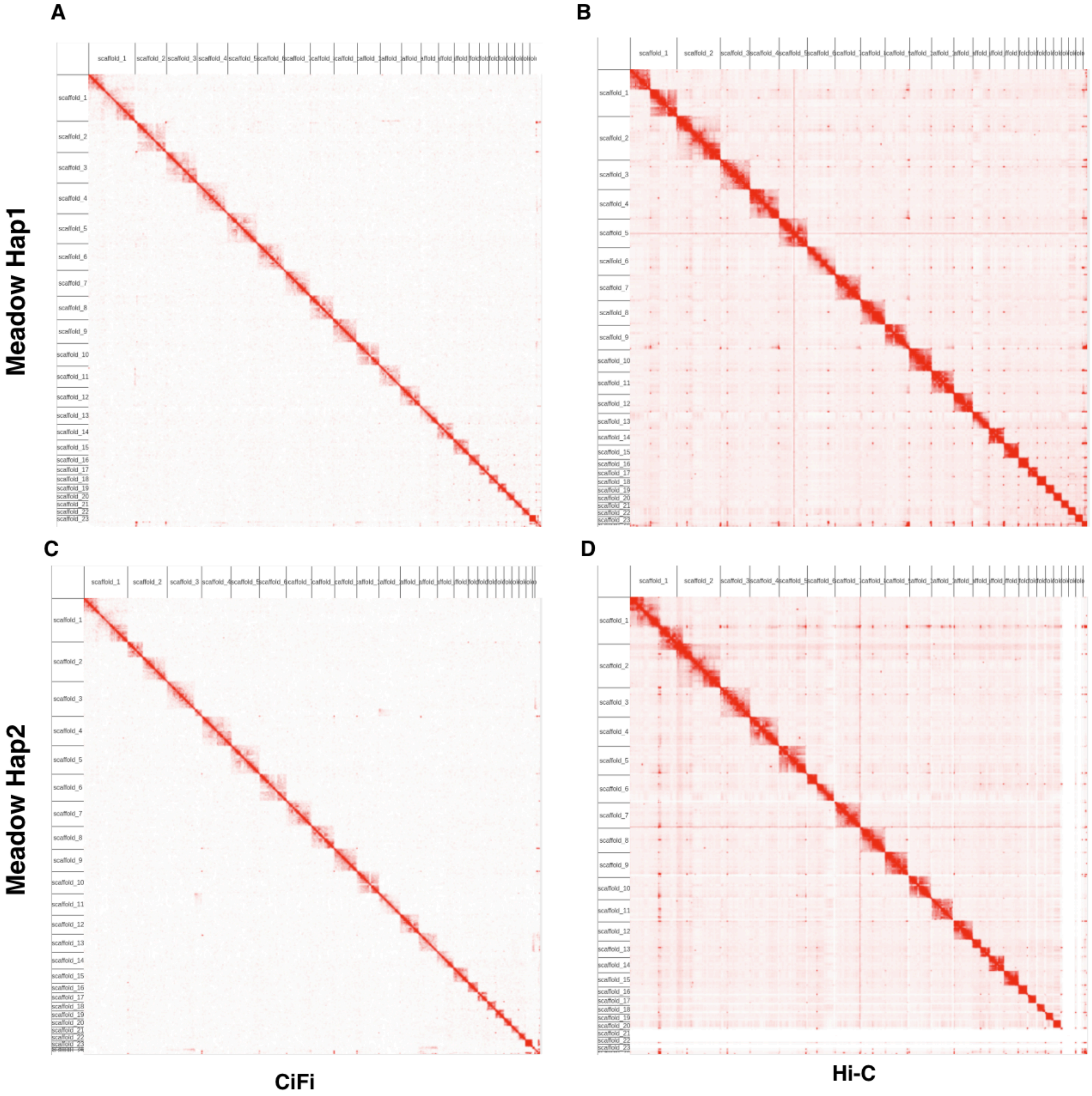

**Figure S2. Comparison of Hi-C and CiFi contact maps for uncurated haplotype-resolved meadow vole genome assemblies, related to Figure 1.** Contact matrices were generated by mapping CiFi (**A**, **C**) and Hi-C (**B**, **D**) reads to YAHS-scaffolded assemblies for both haplotypes. Contact matrices were generated using wf-pore-c and visualized in Juicebox.

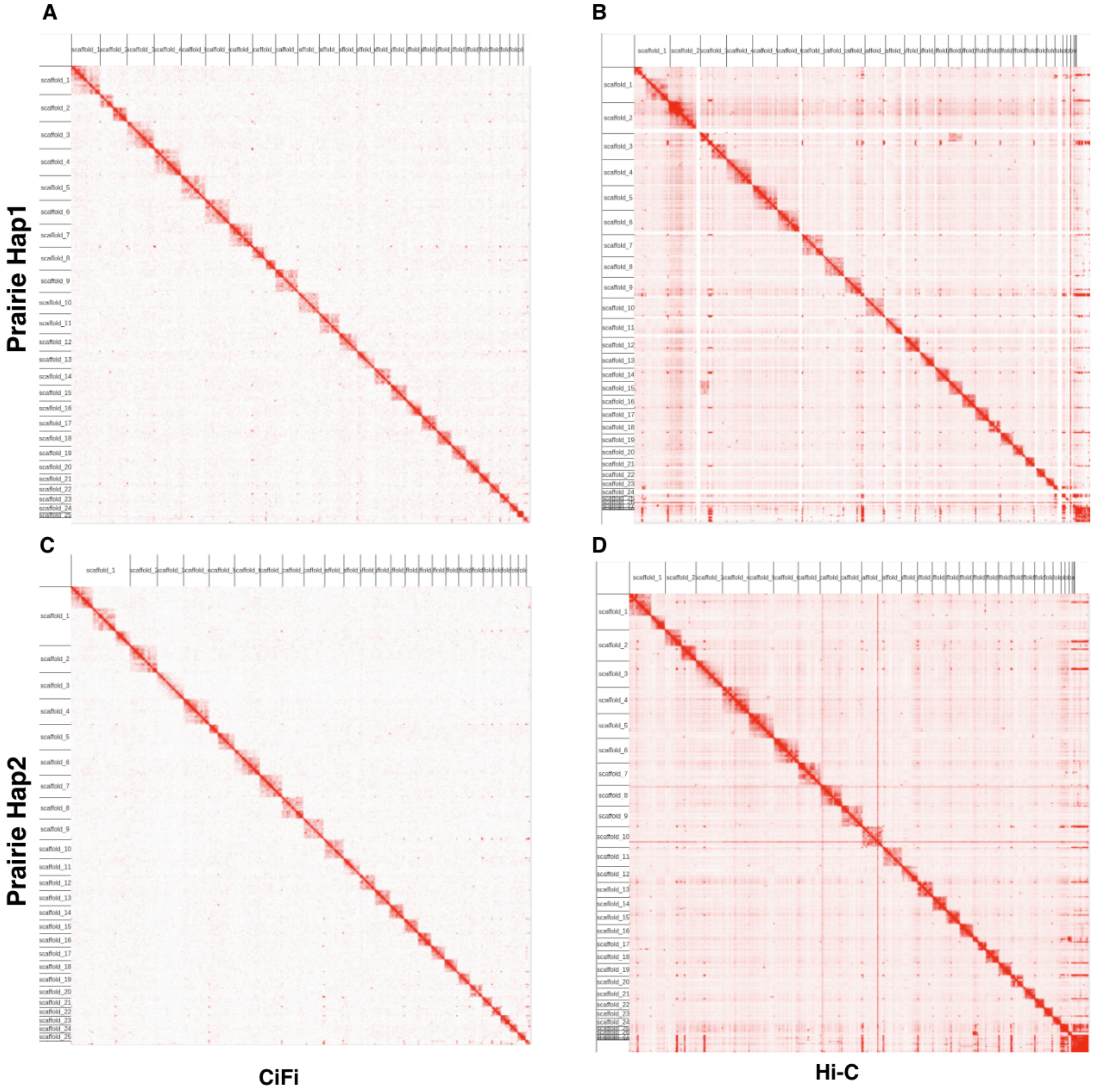

**Figure S3. Comparison of Hi-C and CiFi contact maps for haplotype-resolved uncurated prairie vole genome assemblies, related to Figure 1.** Contact matrices were generated by mapping CiFi (**A**, **C**) and Hi-C (**B**, **D**) reads to YAHS-scaffolded assemblies for both haplotypes. Contact matrices were generated using wf-pore-c and visualized in Juicebox.

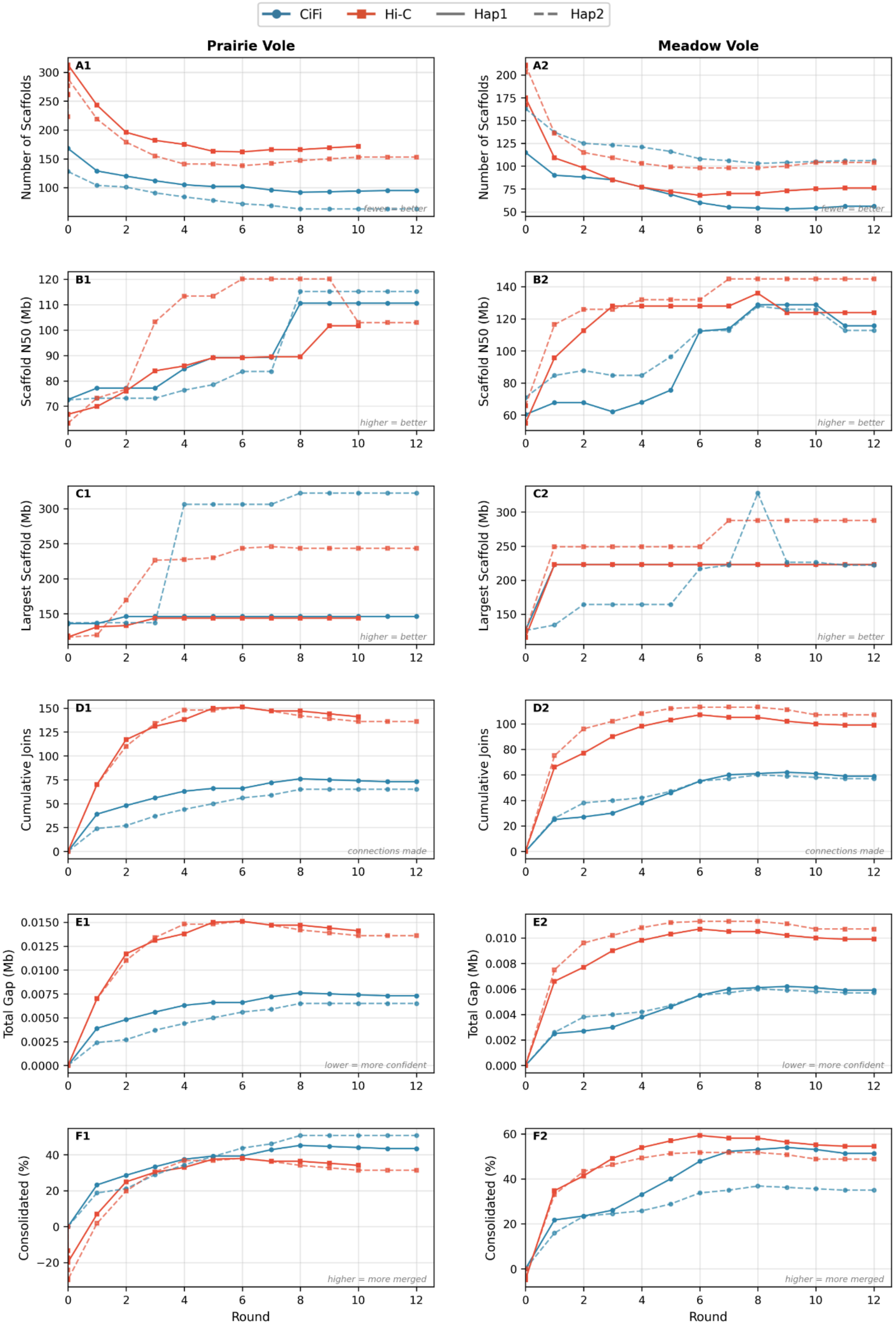

**Figure S4. YAHS scaffolding progression comparing CiFi and Hi-C for prairie vole and meadow vole haplotype-resolved assemblies, related to Figure 1.** Six metrics were tracked across YAHS scaffolding rounds for prairie vole (2n=54) and meadow vole (2n=46) haplotype-resolved assemblies generated by hifiasm dual-scaf, using the same 3C data type for both phasing and downstream scaffolding at matched coverage.

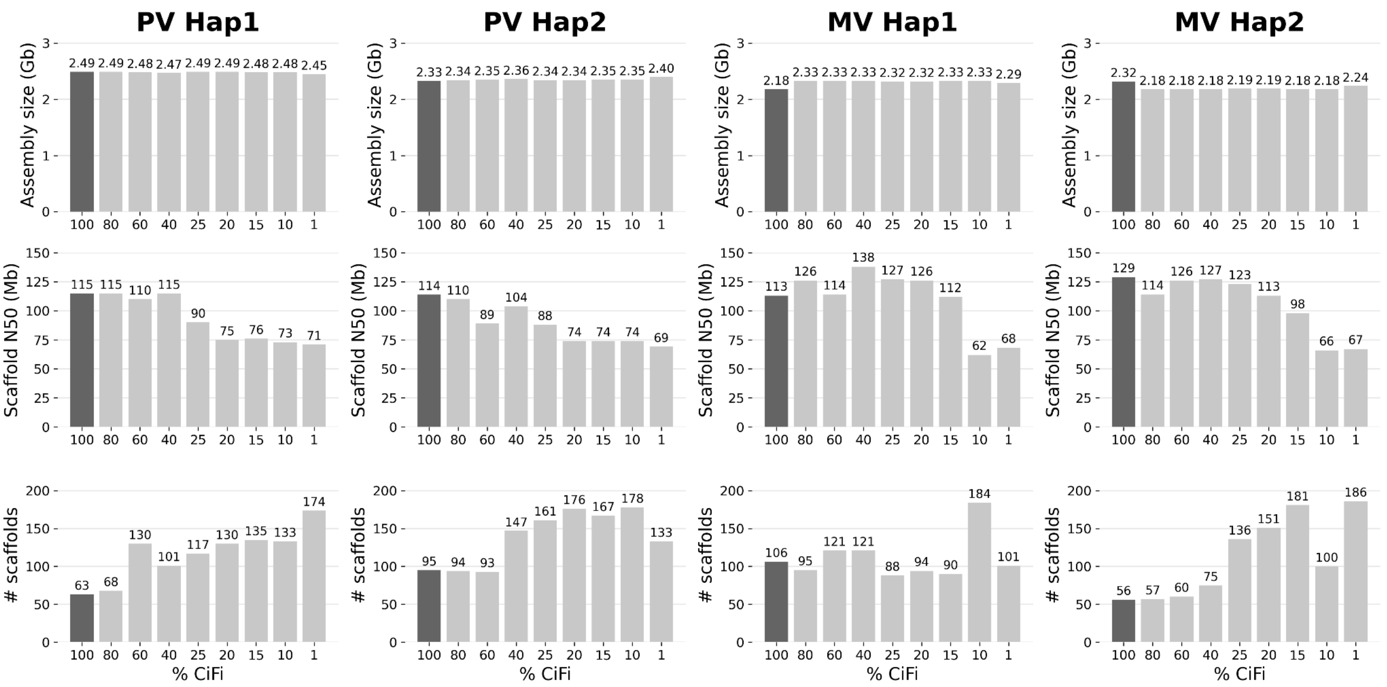

**Figure S5. Effect of CiFi sequencing depth on assembly contiguity, related to Figure 1.** To determine the minimum CiFi input required for chromosome-scale scaffolding, CiFi reads from prairie vole (PV) and meadow vole (MV) were randomly downsampled from 100% to 1% of the original library and used to assemble and scaffold full-depth HiFi assemblies with hifiasm –dual-scaf and YAHS. Assembly size (top), scaffold N50 (middle), and scaffold count (bottom) are shown for each haplotype across all downsampling levels. Assembly size remained stable across all conditions (2.18-2.49 Gb). Scaffold N50 and scaffold count were both sensitive to CiFi depth: all four haplotypes maintained near-optimal N50 values (113–129 Mbp) down to 40% input, with progressive decline at lower levels (66–71 Mbp at 1%). Scaffold counts increased correspondingly at reduced depths, reflecting fragmentation of chromosome-scale scaffolds. These results indicate that ~2× genome coverage of CiFi data is sufficient for high-quality scaffolding, with diminishing returns above 40–60% input. Dark bars indicate the full-depth (100%) condition.

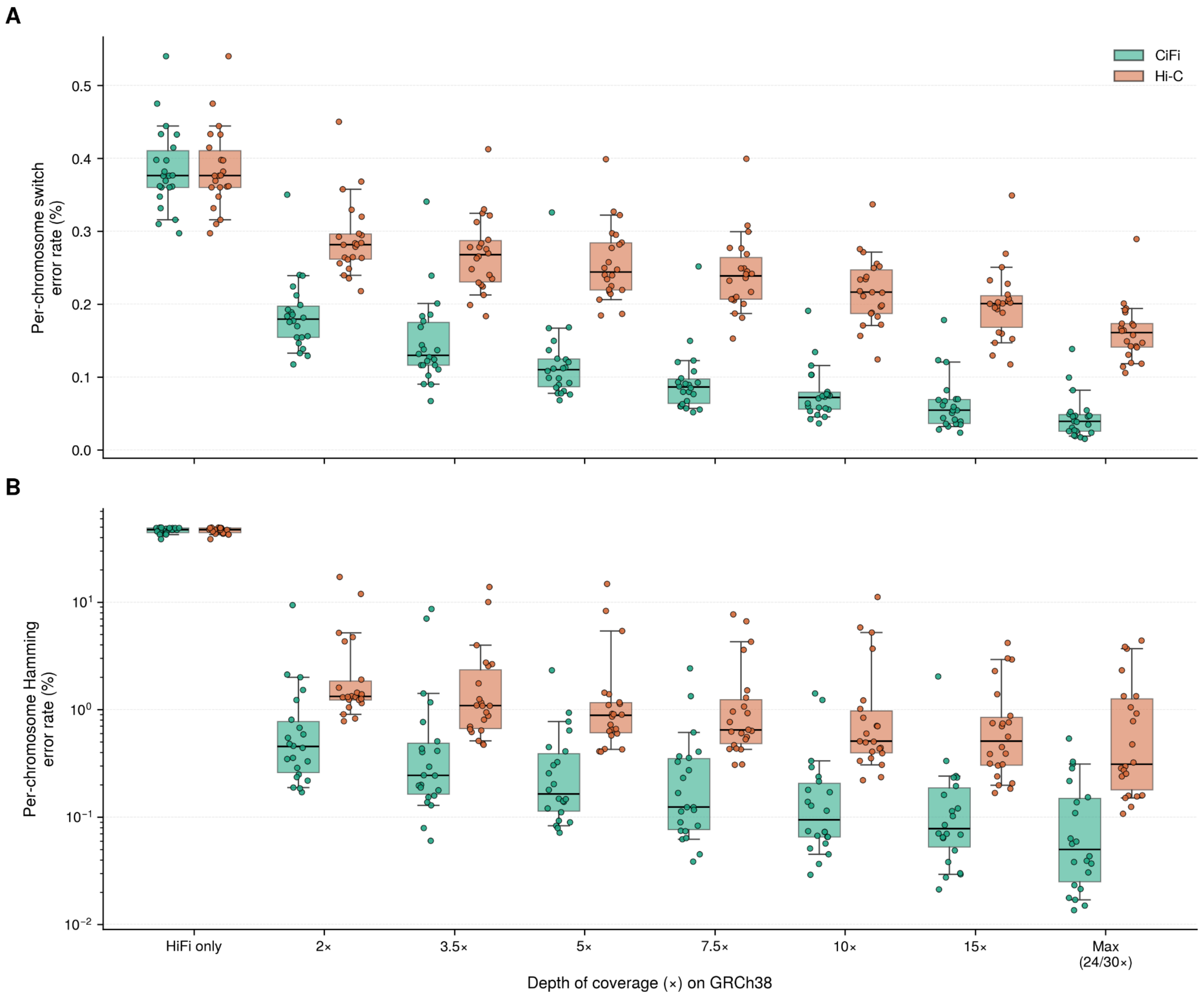

**Figure S6. Phasing accuracy of single-library CiFi versus Hi-C across sequencing depth, related to Figure 1.** Per-chromosome phasing error rates for diploid assemblies of NA12878 (HG001) generated with hifiasm from PacBio HiFi reads alone (HiFi only), or from HiFi reads supplemented with CiFi (teal) or Hi-C (orange) reads, downsampled to the indicated depths on GRCh38. **(A)** Switch error rate and **(B)** blockwise Hamming error rate, shown on a logarithmic scale. The two HiFi-only boxes in each panel represent the same HiFi-only assembly and serve as a shared baseline. "Max" denotes the full available depth of each library (CiFi, 24×; Hi-C, 30×). Each point is one autosome (n = 22; chromosomes 1–22); boxes show the median and interquartile range, and whiskers extend to 1.5× the interquartile range. Diploid assemblies were aligned to GRCh38 with dipcall, and phasing errors were quantified with WhatsHap compared against a trio-phased benchmark phasing of NA12878. Switch error measures local phase flips between consecutive heterozygous variants, whereas blockwise Hamming error measures the minimum fraction of variants assigned to the incorrect benchmark haplotype within phase blocks. At matched downsampled depths, CiFi produces lower error rates than Hi-C by both measures. Median switch error decreases from 0.38% (HiFi only) to 0.04% for CiFi at maximum depth, versus 0.16% for Hi-C. Median Hamming error decreases from 47% (HiFi only) to 0.05% for CiFi at maximum depth, versus 0.31% for Hi-C, indicating that a single CiFi library provides more accurate chromosome-scale phasing than Hi-C at comparable coverage.

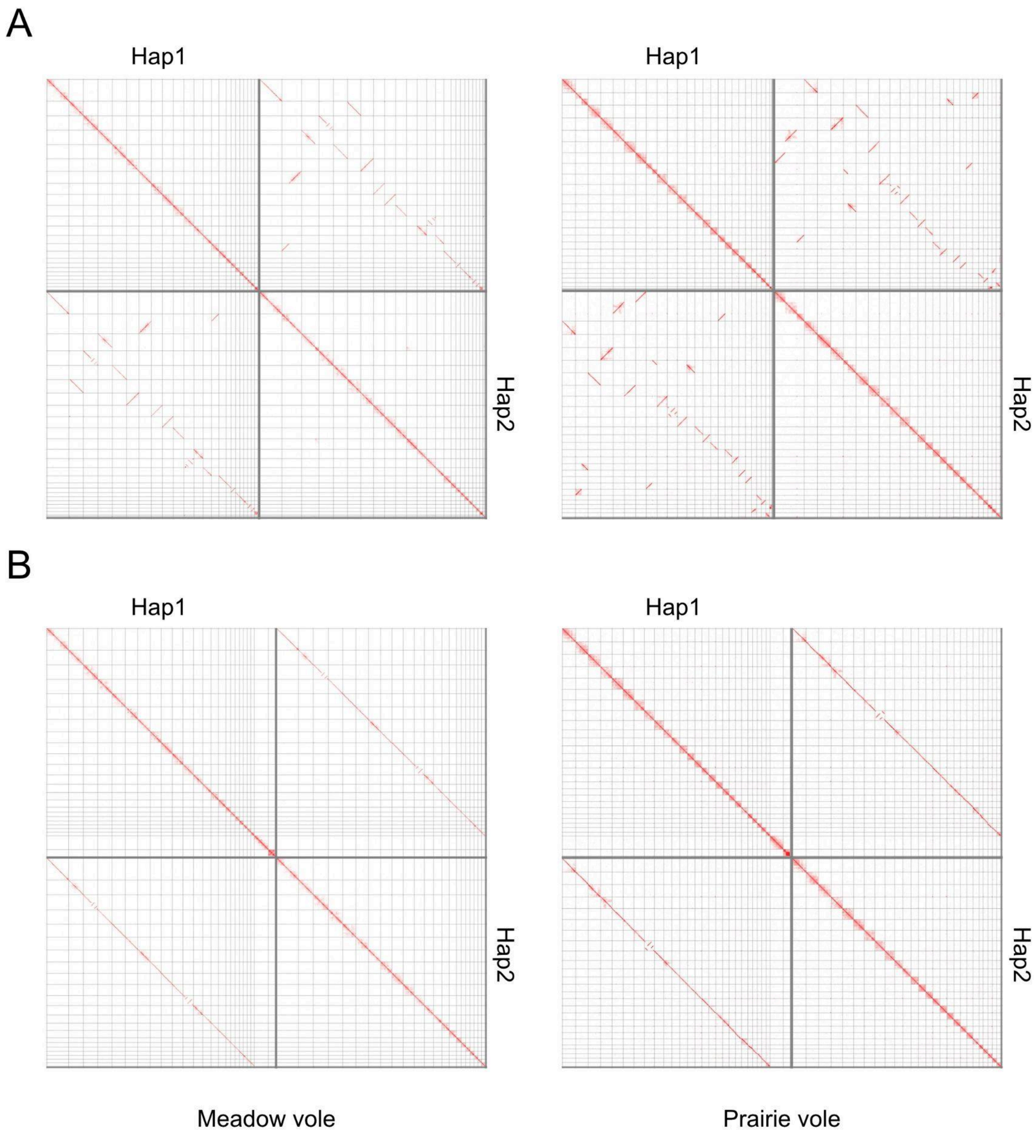

**Figure S7. Curation of prairie and meadow vole assemblies, related to Figure 1.** Contact maps showing the combined haplotype assemblies of the meadow vole and prairie vole **(A)** pre-curation and **(B)** post-curation chromosomally resolved with Hap1 containing chrX and chrY. Contact matrices were generated using wf-pore-c and visualized in HiGlass (see Methods).

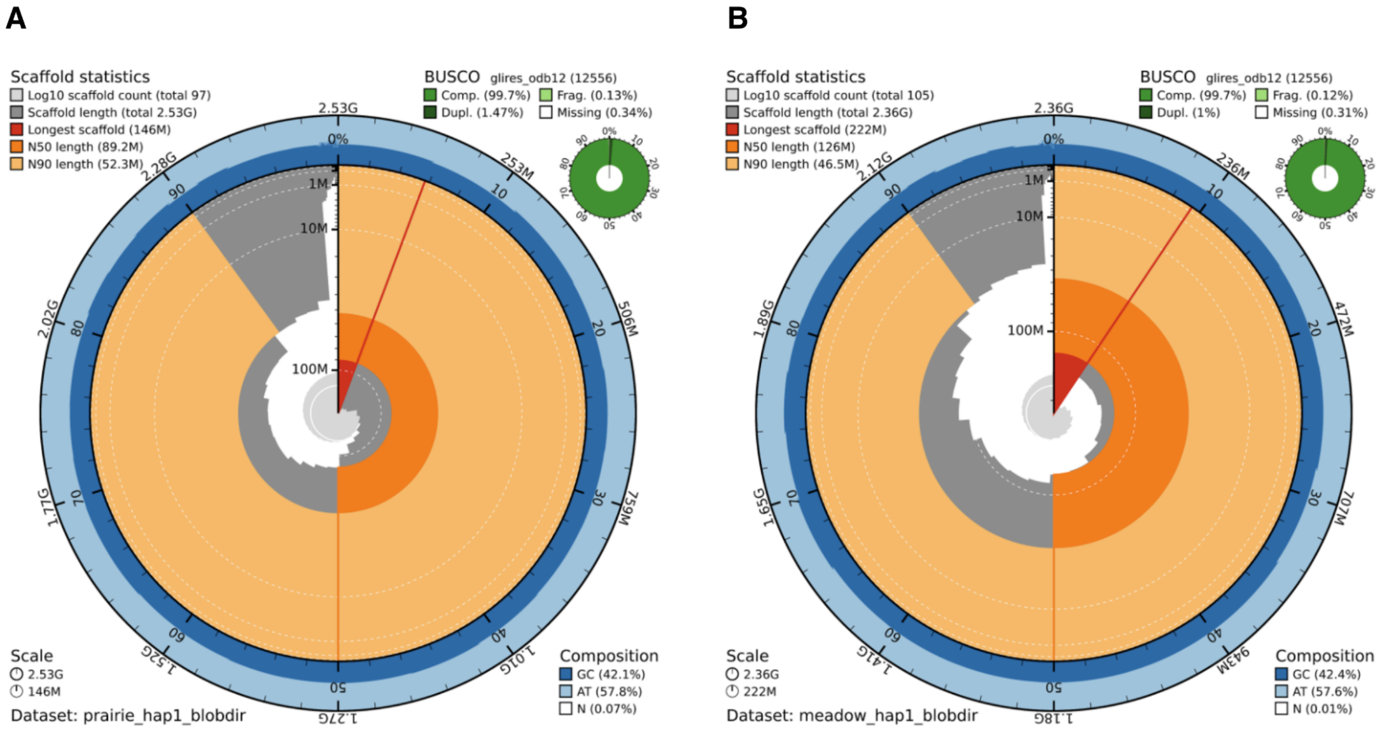

**Figure S8. Assembly metrics for Hap1 vole assemblies, related to Table 1.** BlobToolKit snail plots showing scaffold statistics for prairie (left) and meadow (right) vole Hap1 assemblies. The outer rings display scaffold lengths arranged from longest (red line) to shortest, with cumulative length shown on the inner orange ring. Grey segments indicate scaffold count ($\log_{10}$ scale). Both assemblies show high contiguity, with prairie vole totaling 2.53 Gbp across 97 scaffolds (N50 = 89.2 Mbp) and meadow vole totaling 2.36 Gbp across 105 scaffolds (N50 = 126 Mbp). The outer blue rings represent GC content (~42%) and AT content (~58%). Inset BUSCO plots (glires_odb12, n = 12,556) show near-complete gene coverage (99.7%) for both assemblies, with minimal fragmented or missing orthologs.

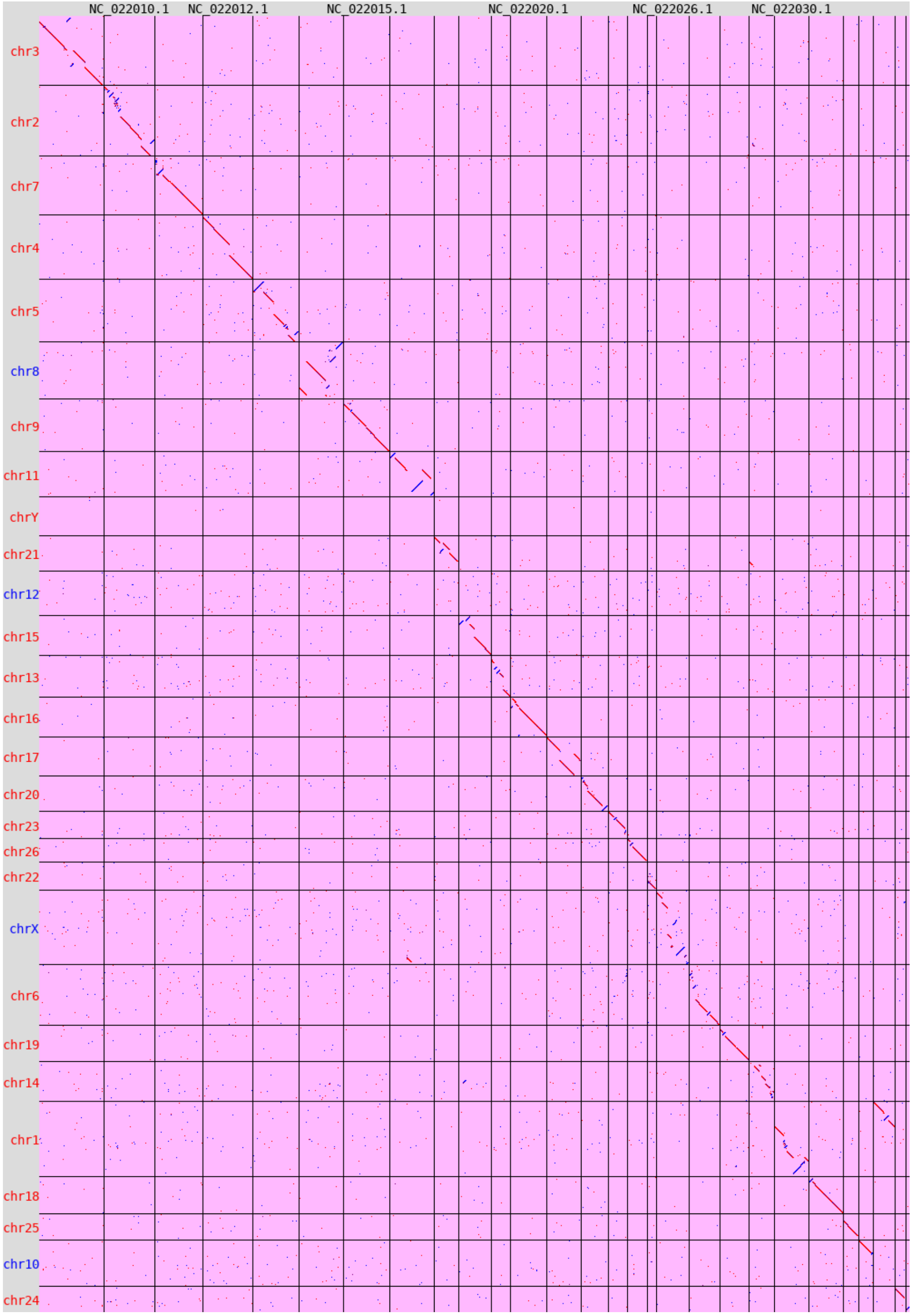

**Figure S9. Comparison of the new prairie vole curated assembly to the short-read based reference genome MicOch1.0, related to Table 1.** Dot plot generated with nf-core/pairgenomealign (see Methods) comparing the new Hap1 curated assembly (y-axis) to MicOch1.0 (x-axis). Each point represents a local pairwise alignment; points along the diagonal indicate collinear, syntenic regions. The near-continuous diagonal confirms broad structural conservation between assemblies, while the fragmented x-axis reflects the substantially lower contiguity of the short-read reference.

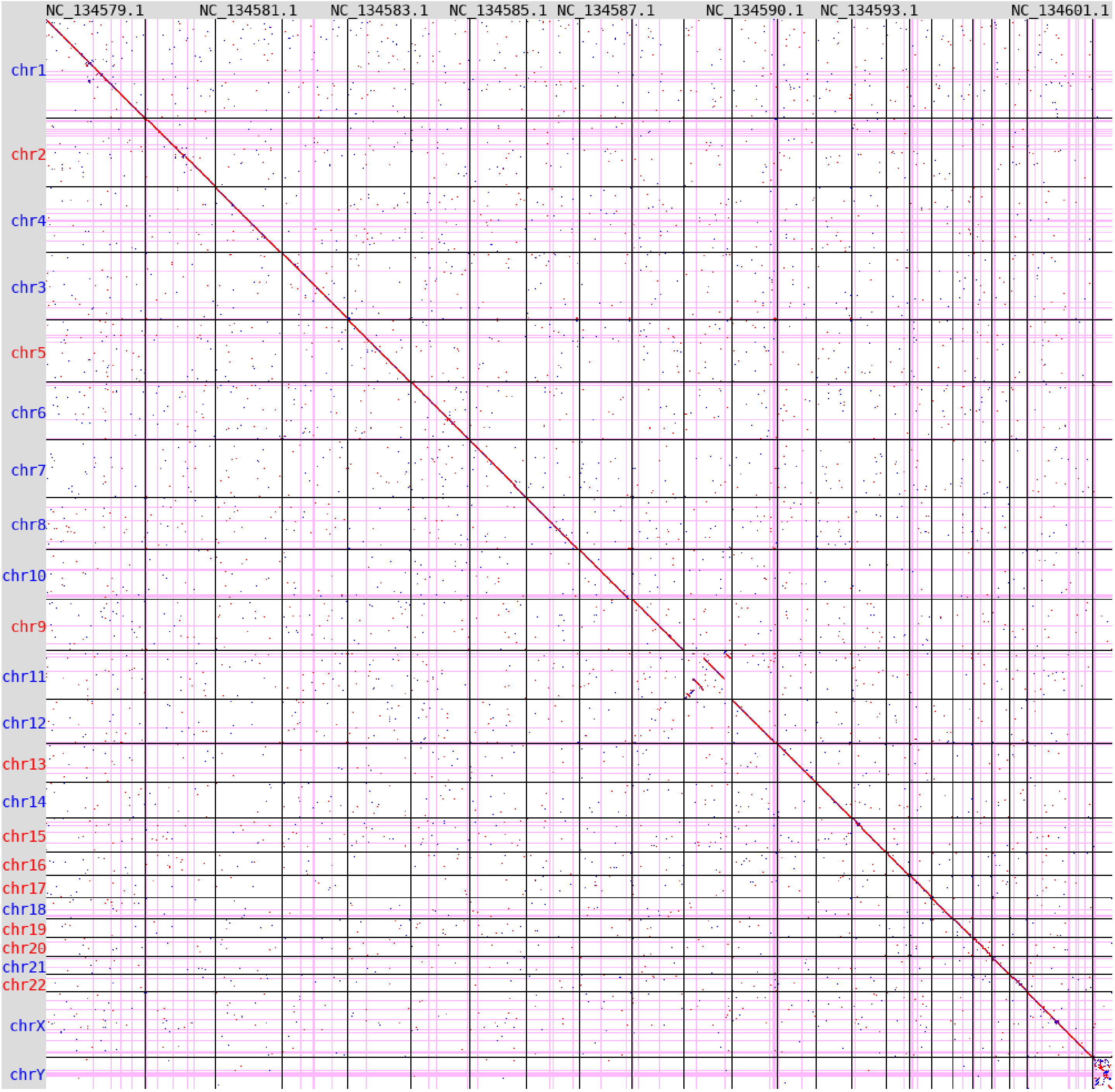

**Figure S10. Comparison of the new meadow vole curated assembly (Hap1) to the VGP (mMicPen1), related to Table 1.** Dot plot, as in Figure S9, comparing the new Hap1 curated assembly (y-axis) to mMicPen1 (x-axis). Each point represents a local pairwise alignment; points along the diagonal indicate collinear, syntenic regions. The near-continuous diagonal confirms high synteny between the two independent long-read assemblies.

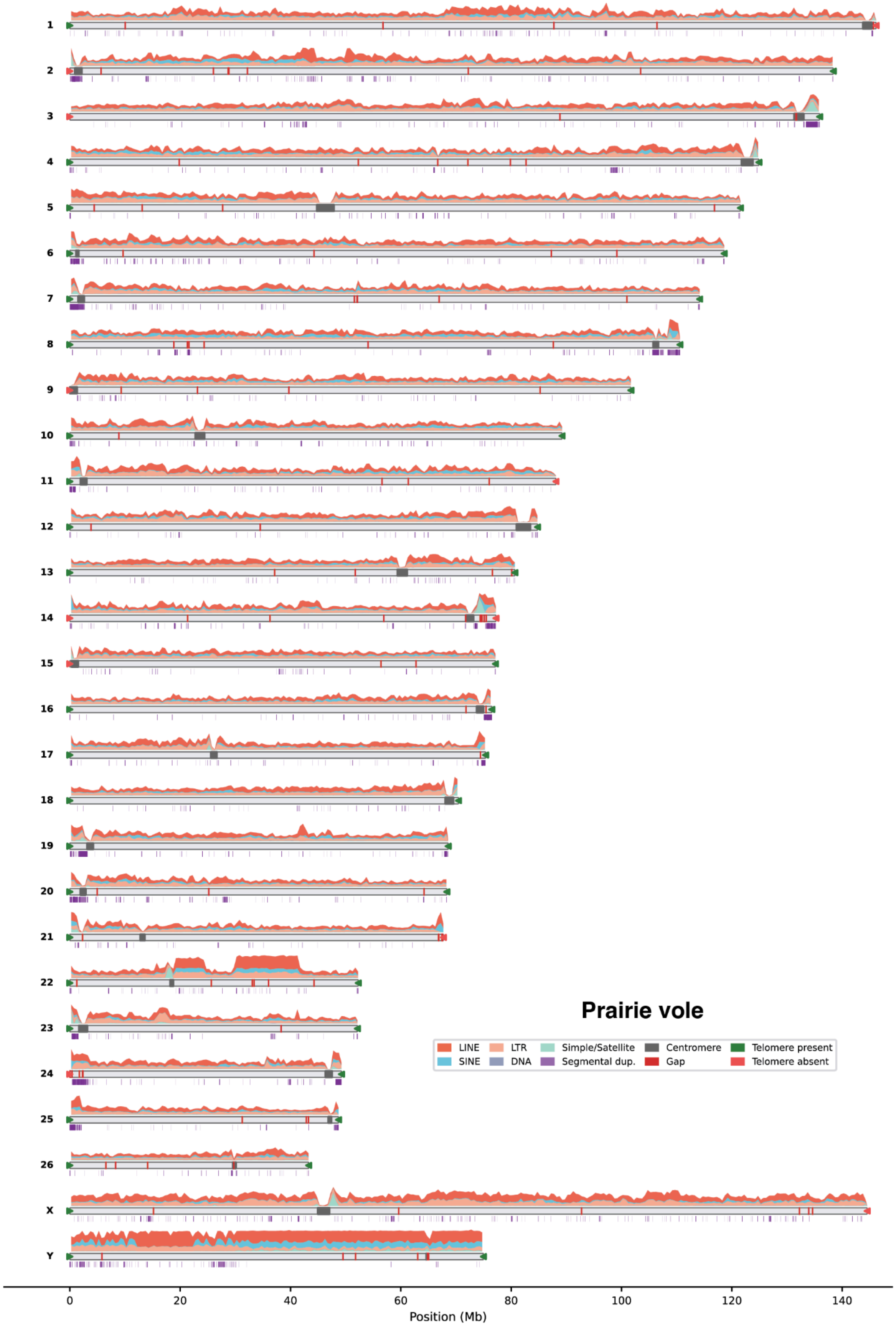

**Figure S11. Annotation of repeat elements and segmental duplications for prairie vole assembly, related to Figure 2.** Repeat and segmental-duplication distributions are shown above and below the Hap1 chromosomal ideograms, respectively.

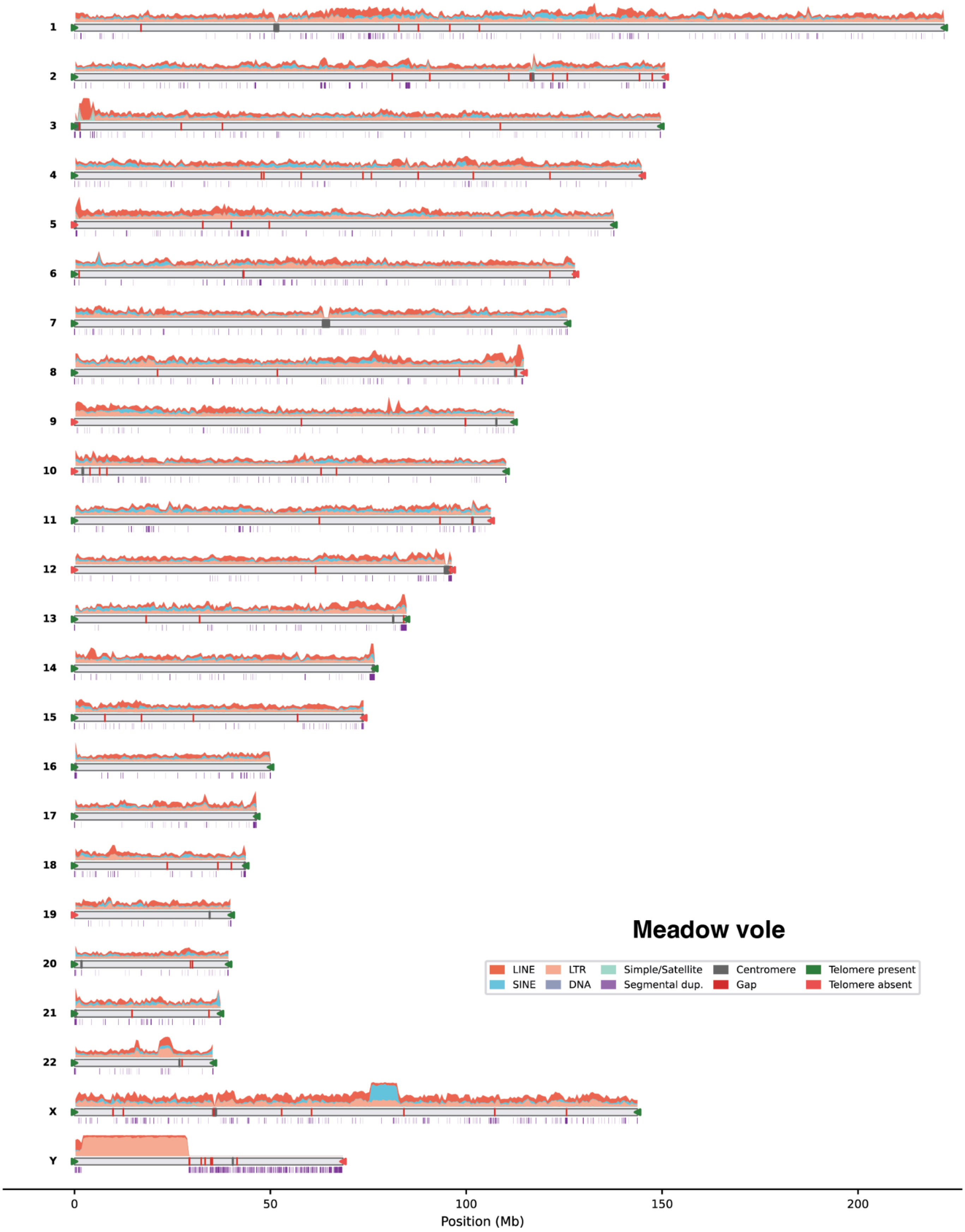

**Figure S12. Annotation of repeat elements and segmental duplications for meadow vole assembly, related to Figure 2.** Repeat and segmental-duplication distributions are shown above and below the Hap1 chromosomal ideograms, respectively.

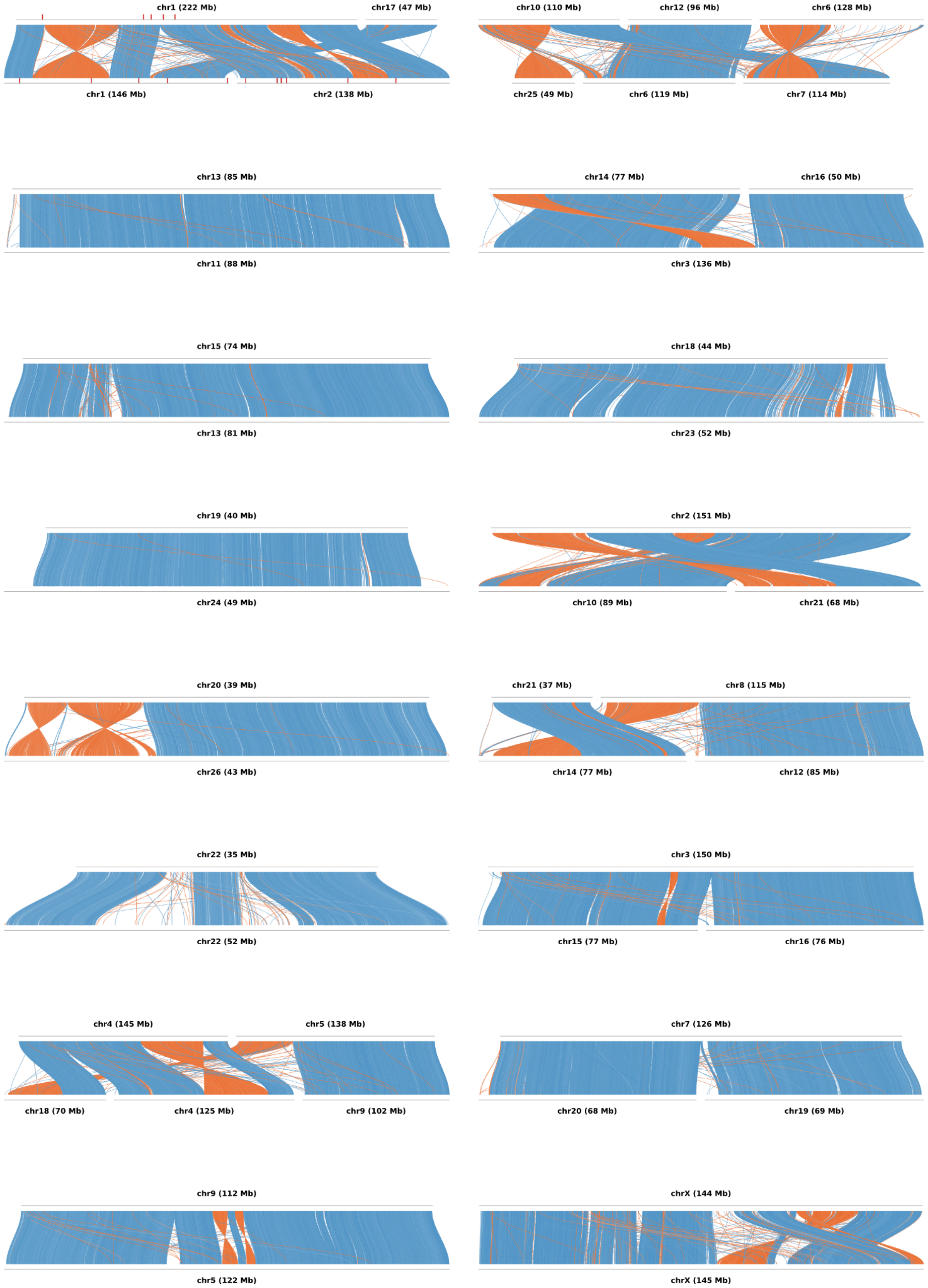

**Figure S13. Chromosome-scale synteny between meadow and prairie vole assemblies, related to Figure 3.** Pairwise MUMmer (nucmer) alignments, visualized with pyGenomeViz, between meadow vole (top track of each panel) and prairie vole (bottom track). Ribbons are coloured blue (forward) or orange (inverted), and red marks on each track denote assembly gaps.

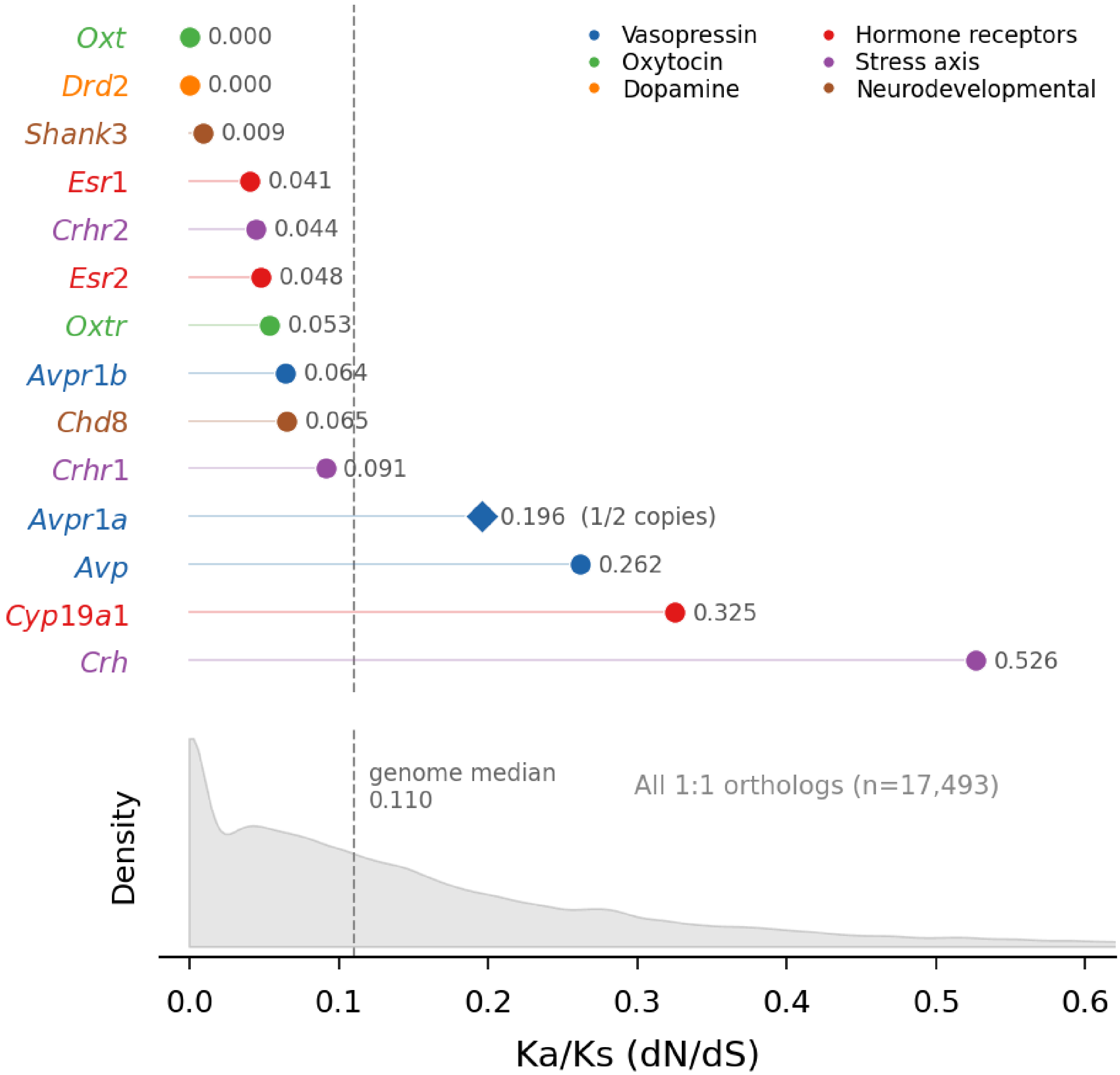

**Figure S14. Ka/Ks landscape of social behavior genes relative to genome-wide distribution, related to Figure 3.** Pairwise meadow–prairie Ka/Ks for 14 manually curated autosomal social behavior genes across six functional pathways, compared with the genome-wide distribution of 17,493 1:1 gene pairs (median Ka/Ks = 0.110). Ka/Ks values were computed in BioPython v1.86 by applying the VEP-annotated SNV substitutions from the meadow-vs-prairie assembly alignment to the prairie EGAPx reference coding sequence and scoring each gene by the Nei–Gojobori (1986) method [1]. For Avpr1a, Ka/Ks was computed between the meadow copy (MPEN1_004149) and prairie copy 1 (MOCH1_002538).

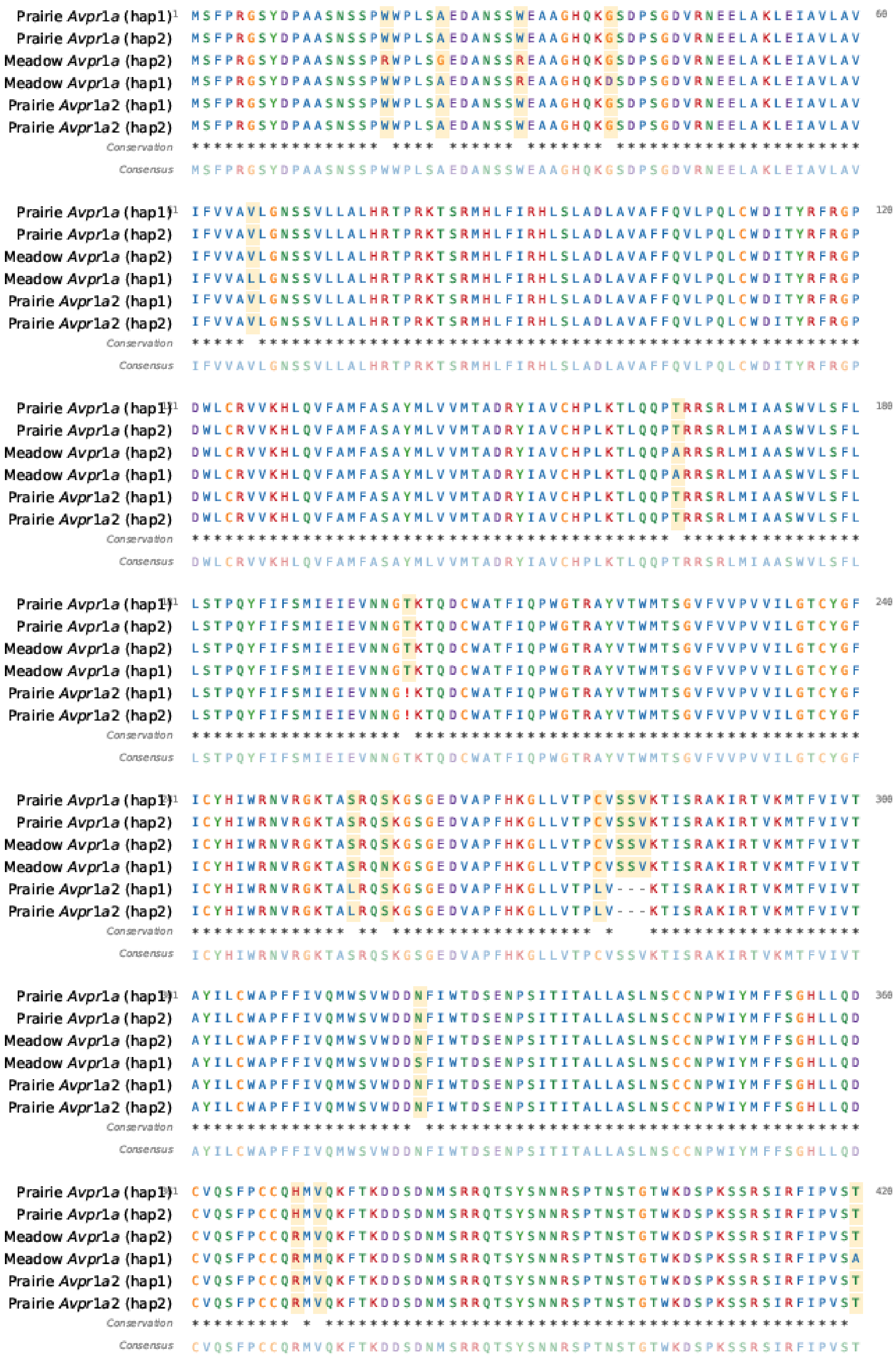

**Figure S15. Amino acid alignment of Avpr1a homologs translated from gene annotations in prairie and meadow vole assemblies, related to Figure 3.** Alignment of both haplotypes of Avpr1a homologous amino translated using gene annotation sequences from prairie and meadow vole assemblies. Positions highlighted in yellow represent sites with differences in amino acids, and the “!” (block 4) designation in Avpr1a2 represents a frame-shift variation resulting in an early stop codon.

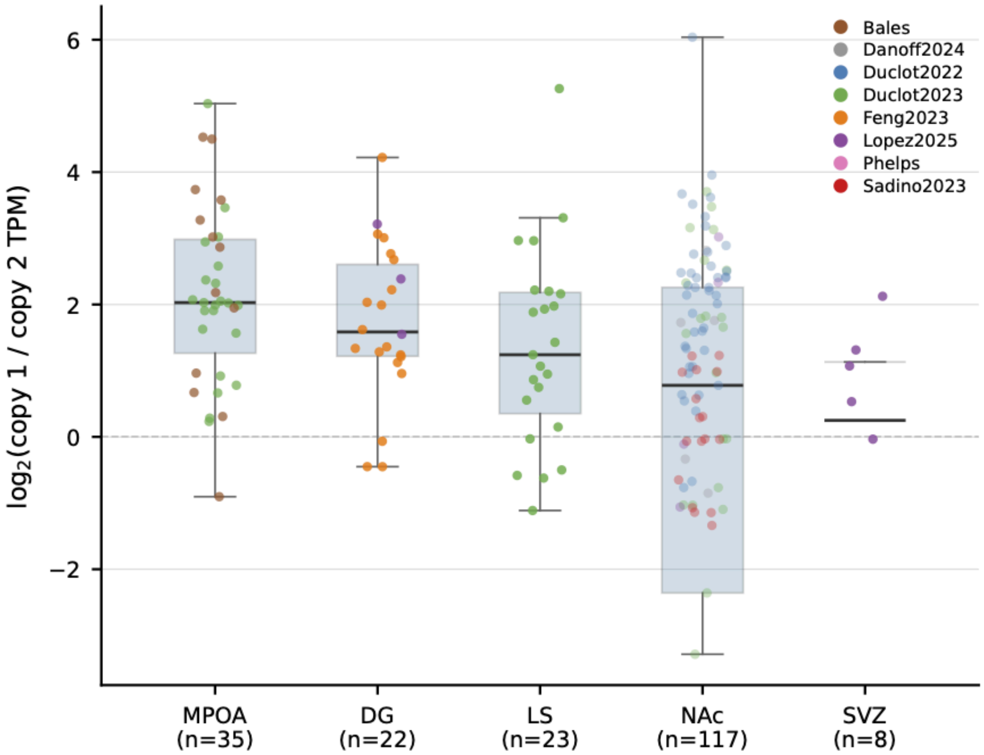

**Figure S16. Transcriptomic analysis of *Avpr1a* paralogs, related to Figure 3.** Transcriptomic analysis of published RNA-seq data [2–7] showing relative expression ($\log_2$ TPM ratio) of *Avpr1a* (copy 1) relative to *Avpr1a2* (copy 2) in medial preoptic area (MPOA), dentate gyrus (DG), lateral septum (LS), nucleus accumbens (NAc), and subventricular zone (SVZ). Points are colored by the study from which RNA-seq data were derived. No expression was detected for either paralog in amygdala, hypothalamus, and ventral pallidum (not shown). TPM, transcripts per million reads.

#### Supplemental Tables

**Table S1. Summary of sequencing metrics, related to Figure 1.**

| Metric | Prairie Vole | Meadow Vole |
|---|---|---|
| Number of reads | 7,216,449 | 7,334,014 |
| Total bases (bp) | 102,662,527,440 | 98,247,171,905 |
| **3C CiFi reads** | | |
| Number of reads | 1,366,318 | 1,544,185 |
| Total bases (bp) | 11,214,456,765 | 11,792,822,905 |
| Average read length (bp) | 8,208 | 7,637 |
| Median read length (bp) | 7,929 | 7,172 |
| Minimum read length (bp) | 337 | 354 |
| Maximum read length (bp) | 21,868 | 22,560 |
| GC content % | 41.6 | 41.9 |
| Total HindIII cut sites | 4,123,365 | 3,587,519 |
| Mean HindIII cut sites per read | 3.0 | 2.3 |
| Median HindIII cut sites per read | 2.0 | 2.0 |
| Reads passing | 1,236,116 | 1,360,368 |
| Pass rate % | 90 | 88 |
| Total fragments | 5,359,481 | 4,947,887 |
| Total pairs | 23,397,997 | 8,120,397 |
| Average fragments per read | 4.3 | 3.6 |
| Average pairs per read | 18.9 | 6.0 |
| Mean fragment size (bp) | 2,043 | 2,298 |
| Median fragment size (bp) | 1,364 | 1,717 |
| Maximum fragment size (bp) | 18,095 | 20,033 |
| mean QV | 44 | 44.14 |
| median QV | 44 | 44.32 |
| PCR duplication read count | 2,465 | 2,062 |
| **DNA HiFi reads** | | |
| Number of reads | 5,662,482 | 5,563,158 |
| Total bases (bp) | 89,452,438,646 | 84,249,860,453 |
| Mean read length (bp) | 15,797 | 15,144 |
| Median read length (bp) | 15,638 | 15,157 |
| Minimum read length (bp) | 71 | 74 |
| Maximum read length (bp) | 55,088 | 50,738 |
| GC content % | 42 | 42 |
| mean QV | 37 | 37 |
| median QV | 36.76 | 36.82 |
| PCR duplication read count | 0 | 0 |

**Table S2. Vole CiFi vs. Hi-C assembly benchmark, related to Figure 1.**

| Species | Haplotype | 3C method | Length (bp) | No. contigs | N50 (bp) | L50 | auN (bp) |
|---|---|---|---|---|---|---|---|
| **Haplotype-resolved contig assembly** | | | | | | | |
| Prairie | 1 | CiFi | 2,494,501,047 | 128 | 44,425,645 | 13 | 74,577,836 |
| | | Hi-C | 2,505,510,878 | 136 | 40,023,958 | 12 | 76,385,652 |
| | 2 | CiFi | 2,334,375,679 | 168 | 39,215,499 | 12 | 73,441,114 |
| | | Hi-C | 2,327,235,065 | 174 | 46,893,501 | 12 | 74,484,558 |
| Meadow | 1 | CiFi | 2,323,597,643 | 115 | 42,190,245 | 13 | 66,012,564 |
| | | Hi-C | 2,336,962,076 | 177 | 42,190,220 | 13 | 64,869,002 |
| | 2 | CiFi | 2,184,589,562 | 163 | 46,507,169 | 13 | 67,117,100 |
| | | Hi-C | 2,170,880,991 | 107 | 46,507,169 | 13 | 66,214,157 |
| **Scaffolded assembly** | | | | | | | |
| Prairie | 1 | CiFi | 2,494,507,547 | 63 | 115,203,824 | 8 | 136,946,035 |
| | | Hi-C | 2,505,526,278 | 171 | 113,504,764 | 8 | 115,509,575 |
| | 2 | CiFi | 2,334,382,979 | 95 | 110,586,647 | 9 | 103,880,896 |
| | | Hi-C | 2,327,249,465 | 155 | 103,466,726 | 8 | 114,964,499 |
| Meadow | 1 | CiFi | 2,323,603,543 | 56 | 115,622,006 | 7 | 128,204,822 |
| | | Hi-C | 2,336,975,176 | 119 | 128,746,723 | 7 | 133,583,981 |
| | 2 | CiFi | 2,184,595,262 | 106 | 112,728,876 | 8 | 112,178,339 |
| | | Hi-C | 2,170,890,091 | 62 | 127,793,347 | 7 | 133,015,360 |

**Table S4. BUSCO genome completeness of curated vole assemblies, related to Table 1.**

| Assembly | Complete (%) | Single-copy (%) | Duplicate (%) | Fragment (%) | Missing (%) | Complete (n) | Single -copy (n) | Duplicate (n) | Fragment (n) | Missing (n) | Total length (bp) |
|---|---|---|---|---|---|---|---|---|---|---|---|
| Prairie vole hap1 (this study) | 99.70 | 98.20 | 1.50 | 0.10 | 0.20 | 12,513 | 12,329 | 184 | 16 | 27 | 2,530,070,934 |
| Prairie vole hap2 (this study) | 97.20 | 95.80 | 1.40 | 0.20 | 2.60 | 12,206 | 12,024 | 182 | 20 | 330 | 2,298,544,460 |
| Meadow vole hap1 (this study) | 99.70 | 98.70 | 1.00 | 0.10 | 0.20 | 12,517 | 12,391 | 126 | 15 | 24 | 2,357,820,369 |
| Meadow vole hap2 (this study) | 97.00 | 96.10 | 0.90 | 0.10 | 2.90 | 12,174 | 12,065 | 109 | 18 | 364 | 2,150,379,936 |
| Prairie vole MicOch1.0 (NCBI reference) | 98.90 | 98.10 | 0.80 | 0.50 | 0.50 | 12,423 | 12,317 | 106 | 65 | 68 | 2,287,340,943 |
| Meadow vole VGP hap1 (mMicPen1) | 99.70 | 98.60 | 1.10 | 0.10 | 0.20 | 12,522 | 12,385 | 137 | 14 | 20 | 2,369,119,668 |
| Meadow vole VGP hap2 (mMicPen1) | 97.10 | 96.20 | 0.90 | 0.10 | 2.70 | 12,194 | 12,081 | 113 | 17 | 345 | 2,156,400,253 |

**Table S6. Prairie vole linkage group validation, related to Table 1.**

| Scaffold | Assigned Chr | Linkage Group | n Markers | Spearman ρ | Linkage Map Call* | Alignment Call* | Alignment Confidence | Concordant |
|---|---|---|---|---|---|---|---|---|
| SUPER_1 | chr1 | LGLG4 | 18 | 0.79 | keep | keep | low_mixed | Yes |
| SUPER_1 | chr1 | LGLG9 | 7 | 0.991 | keep | keep | low_mixed | Yes |
| SUPER_2 | chr2 | LG2 | 21 | 0.998 | keep | keep | low_mixed | Yes |
| SUPER_3 | chr3 | LG1 | 28 | -0.971 | RC | RC | medium | Yes |
| SUPER_4 | chr4 | LG5 | 18 | -0.965 | RC | RC | high | Yes |
| SUPER_5 | chr5 | LG6 | 24 | 0.944 | keep | keep | low_mixed | Yes |
| SUPER_6 | chr6 | LGLG1 | 16 | 0.993 | keep | keep | low_mixed | Yes |
| SUPER_7 | chr7 | LG4 | 19 | 0.933 | keep | keep | medium | Yes |
| SUPER_8 | chr8 | LG7 | 13 | -0.93 | RC | RC | low_mixed | Yes |
| SUPER_9 | chr9 | LG8 | 22 | 0.987 | keep | keep | high | Yes |
| SUPER_10 | chr10 | LG14 | 11 | 0.406 | keep | keep | low_mixed | Yes |
| SUPER_10 | chr10 | LGLG8 | 5 | -0.975 | RC | keep | low_mixed | No |
| SUPER_11 | chr11 | LG10 | 14 | 0.977 | keep | keep | low_mixed | Yes |
| SUPER_12 | chr12 | LG14.2 | 7 | -0.929 | RC | RC | high | Yes |
| SUPER_12 | chr12 | LG14.3 | 5 | -0.1 | RC | RC | high | Yes |
| SUPER_13 | chr13 | LG17 | 13 | -0.995 | RC | RC | low_mixed | Yes |
| SUPER_14 | chr14 | LGLG3 | 13 | -0.899 | RC | RC | medium | Yes |
| SUPER_15 | chr15 | LG16 | 20 | 0.958 | keep | keep | low_mixed | Yes |
| SUPER_16 | chr16 | LG18 | 14 | -0.974 | RC | RC | medium | Yes |
| SUPER_17 | chr17 | LG19 | 12 | 0.845 | keep | keep | high | Yes |
| SUPER_18 | chr18 | LGLG5 | 14 | -0.957 | RC | RC | medium | Yes |
| SUPER_19 | chr19 | LGLG2 | 11 | 0.989 | keep | keep | medium | Yes |
| SUPER_20 | chr20 | LG21 | 12 | 0.993 | keep | keep | low_mixed | Yes |
| SUPER_21 | chr21 | LG15 | 13 | 0.912 | keep | keep | medium | Yes |
| SUPER_23 | chr23 | LG22 | 8 | -0.826 | RC | RC | medium | Yes |
| SUPER_24 | chr24 | LGLG10 | 4 | 0.949 | keep | keep | high | Yes |
| SUPER_25 | chr25 | LGLG7 | 4 | -0.4 | RC | RC | high | Yes |
| SUPER_26 | chr26 | LG24 | 8 | -1 | RC | RC | medium | Yes |
| SUPER_X | chrX | LGX | 11 | 0.596 | keep | RC | low_mixed | No |

* RC, reverse complement

**Table S8. Gene annotations, related to Figure 2.**

| Metric | Prairie vole (this study) | Prairie vole (MicOch1.0) | Meadow vole (this study) | Meadow vole (mMicPen1) |
|---|---|---|---|---|
| **Haplotype** | **Hap1*** | **merged** | **Hap1*** | **Hap1*** |
| **Protein-coding genes** | 20,526 | 20,077 | 20,513 | 21,461 |
| **mRNA transcripts** | 44,342 | 37,795 | 49,821 | 51,588 |
| **Pseudogenes** | 4,521 | 3,794 | 3,719 | 4,163 |
| **lncRNA genes** | 1,060 | 655 | 2,656 | 1,652 |
| **Mean CDS length (bp)** | 2,108 | 2,002 | 2,156 | 2,149 |
| **Overlapping genes** | 598 | 1,412 | 771 | 1,795 |

* Gene annotation was performed and compared only for primary Hap1 assemblies

**Table S9. Repeat annotations, related to Figure 2.**

| Repeat Type | Prairie Count | Prairie masked (bp) | Prairie % | Meadow Count | Meadow Masked (bp) | Meadow % |
|---|---|---|---|---|---|---|
| DNA | 0 | 0 | 0.00% | 583 | 79,788 | 0.00% |
| Kolobok-T2 | 0 | 0 | 0.00% | 1,436 | 255,187 | 0.01% |
| TcMar-Mariner | 0 | 0 | 0.00% | 167 | 70,255 | 0.00% |
| TcMar-Tigger | 16,061 | 2,866,823 | 0.11% | 16,194 | 2,768,807 | 0.12% |
| hAT | 343 | 50,911 | 0.00% | 339 | 49,708 | 0.00% |
| hAT-Blackjack | 824 | 94,126 | 0.00% | 870 | 98,219 | 0.00% |
| hAT-Charlie | 47,018 | 8,522,287 | 0.34% | 45,690 | 8,756,479 | 0.37% |
| hAT-Tip100 | 344 | 42,961 | 0.00% | 0 | 0 | 0.00% |
| L1 | 1,532,863 | 422,944,831 | 16.72% | 1,046,072 | 306,670,347 | 13.01% |
| L1-Tx1 | 2,072 | 896,117 | 0.04% | 1,622 | 770,793 | 0.03% |
| L2 | 1,623 | 93,820 | 0.00% | 1,007 | 72,019 | 0.00% |
| RTE-BovB | 818 | 433,541 | 0.02% | 952 | 510,900 | 0.02% |
| I | 4,051 | 2,386,343 | 0.09% | 0 | 0 | 0.00% |
| LTR | 1,979 | 1,054,645 | 0.04% | 2,957 | 1,415,059 | 0.06% |
| ERV | 0 | 0 | 0.00% | 1,866 | 674,817 | 0.03% |
| ERV1 | 185,777 | 38,861,168 | 1.54% | 79,349 | 25,929,729 | 1.10% |
| ERVK | 547,487 | 213,458,329 | 8.44% | 563,311 | 183,117,344 | 7.77% |
| ERVL | 77,933 | 14,452,654 | 0.57% | 47,824 | 12,551,523 | 0.53% |
| ERVL-MaLR | 232,025 | 59,803,882 | 2.36% | 264,249 | 63,562,566 | 2.70% |
| Copia | 1,861 | 382,438 | 0.02% | 0 | 0 | 0.00% |
| Gypsy | 4,374 | 811,531 | 0.03% | 0 | 0 | 0.00% |
| L1-dep | 21,865 | 7,537,878 | 0.30% | 22,316 | 7,899,150 | 0.34% |
| 5S | 2,885 | 669,903 | 0.03% | 4,462 | 531,407 | 0.02% |
| Alu | 661,291 | 78,256,062 | 3.09% | 653,454 | 86,587,172 | 3.67% |
| B2 | 274,718 | 39,738,970 | 1.57% | 376,973 | 46,065,396 | 1.95% |
| B4 | 35,254 | 3,749,306 | 0.15% | 15,455 | 2,083,391 | 0.09% |
| ID | 111,286 | 19,544,828 | 0.77% | 15,255 | 956,945 | 0.04% |
| MIR | 2,403 | 191,797 | 0.01% | 6,863 | 533,669 | 0.02% |
| U | 0 | 0 | 0.00% | 258 | 19,805 | 0.00% |
| tRNA | 22,867 | 2,325,496 | 0.09% | 23,194 | 2,095,772 | 0.09% |
| tRNA-RTE | 200 | 22,648 | 0.00% | 212 | 24,688 | 0.00% |
| Unknown | 58,691 | 57,166,588 | 2.26% | 52,654 | 12,293,048 | 0.52% |
| **Total interspersed** | **3,848,913** | **976,359,883** | **38.59%** | **3,245,584** | **766,443,983** | **32.51%** |
| Low_complexity | 89,018 | 4,932,896 | 0.19% | 91,176 | 5,313,993 | 0.23% |
| Satellite | 9,020 | 8,617,311 | 0.34% | 2,716 | 511,150 | 0.02% |
| Simple_repeat | 809,325 | 40,301,928 | 1.59% | 704,738 | 36,378,666 | 1.54% |
| rRNA | 480 | 1,691,660 | 0.07% | 11,120 | 1,987,457 | 0.08% |
| snRNA | 2,161 | 255,051 | 0.01% | 1,945 | 237,980 | 0.01% |
| **Total** | **4,758,917** | **1,032,158,729** | **40.80%** | **4,057,279** | **810,873,229** | **34.39%** |

**Table S10. Segmental duplications (SDs) annotations, related to Figure 2.**

| Metric | Meadow vole | Prairie vole |
|---|---|---|
| **Overview** | | |
| Total SD pairs [1] | 5,266 | 5,281 |
| Total SD coverage (merged, non-redundant) [2] | 28.07 Mbp | 54.01 Mbp |
| Total SD bases (sum of pair averages) [3] | 43.26 Mbp | 80.03 Mbp |
| Chromosomes with SDs | 24 | 28 |
| **Gene Overlap** | | |
| SD bp overlapping ≥1 gene | 20.19 Mb | 42.45 Mb |
| SD bp with no gene overlap | 10.93 Mb | 16.34 Mb |
| % of SD bp overlapping genes | 72.40% | 78.90% |
| Intra-chromosomal SD bp (merged) | 21.59 Mb | 24.01 Mb |
| Inter-chromosomal SD bp (merged) | 9.62 Mb | 34.56 Mb |
| Genes overlapping SDs | 1,194 | 1,151 |
| Total gene-body bp inside SDs | 8.98 Mbp | 10.43 Mbp |
| Mean SD region size (gene-overlapping) | 11.7 kb | 23.3 kb |
| Median SD region size (gene-overlapping) | 3.0 kb | 3.9 kb |
| **Chromosomal Distribution** | | |
| Intra-chromosomal pairs [4] | 2,827 | 1,906 |
| Intra-chromosomal coverage (merged) [5] | 21.69 Mbp | 24.11 Mbp |
| Intra-chromosomal bases (sum of pair avgs) [6] | 27.86 Mbp | 25.09 Mbp |
| Inter-chromosomal pairs [7] | 2,439 | 3,375 |
| Inter-chromosomal coverage (merged) [8] | 9.74 Mbp | 34.67 Mbp |
| Inter-chromosomal bases (sum of pair avgs) [9] | 15.40 Mbp | 54.95 Mbp |
| **Size Statistics** | | |
| Mean size (bp) | 8,214.55 | 15,155.15 |
| Median size (bp) | 3,018.25 | 3,104.50 |
| Min size (bp) | 1,000.50 | 1,000 |
| Max size (bp) | 526,368.50 | 1,717,934.50 |
| **Identity Statistics** | | |
| Mean identity (%) | 93.69% | 94.21% |
| Median identity (%) | 93.40% | 94.00% |
| Min identity (%) | 90.00% | 90.00% |
| Max identity (%) | 100.00% | 100.00% |
| **Identity Thresholds (pairs / % of total / bases)** | | |
| >90% identity – pairs | 51.93 | 52.05 |
| >90% identity – % of total | 0.99 | 0.99 |
| >90% identity – bases (bp) | 429,757.62 | 793,243.51 |
| >91% identity – pairs | 43.44 | 46.11 |
| >91% identity – % of total | 0.82 | 0.87 |
| >91% identity – bases (bp) | 375,516.52 | 698,528.36 |
| >92% identity – pairs | 36.88 | 39.42 |
| >92% identity – % of total | 0.70 | 0.75 |
| >92% identity – bases (bp) | 334,619.04 | 581,602.26 |
| >93% identity – pairs | 28.71 | 33.15 |

| | | |
|---|---|---|
| >93% identity – % of total | 0.55 | 0.63 |
| >93% identity – bases (bp) | 277,578.59 | 460,192.42 |
| >94% identity – pairs | 21.47 | 26.31 |
| >94% identity – % of total | 0.41 | 0.50 |
| >94% identity – bases (bp) | 210,821.97 | 338,341.13 |
| >95% identity – pairs | 15.16 | 19.66 |
| >95% identity – % of total | 0.29 | 0.37 |
| >95% identity – bases (bp) | 155,464.23 | 268,400.24 |
| >96% identity – pairs | 10.51 | 14.57 |
| >96% identity – % of total | 0.20 | 0.28 |
| >96% identity – bases (bp) | 110,993.80 | 172,197.38 |
| >97% identity – pairs | 6.71 | 8.66 |
| >97% identity – % of total | 0.13 | 0.16 |
| >97% identity – bases (bp) | 70,098.46 | 104,614.43 |
| >98% identity – pairs | 2.54 | 4.21 |
| >98% identity – % of total | 0.05 | 0.08 |
| >98% identity – bases (bp) | 22,992.49 | 36,484.64 |
| >99% identity – pairs | 0.94 | 1.82 |
| >99% identity – % of total | 0.02 | 0.03 |
| >99% identity – bases (bp) | 9,706.95 | 17,688.21 |
| **Size Distribution (pairs / % of total / bases)** | | |
| 1–5 kb – pairs | 3,825 | 3,301 |
| 1–5 kb – % of total | 72.64% | 62.51% |
| 1–5 kb – bases (bp) | 9,061,440 | 6,900,325 |
| 5–10 kb – pairs | 769 | 869 |
| 5–10 kb – % of total | 14.60% | 16.46% |
| 5–10 kb – bases (bp) | 5,351,702 | 5,932,098 |
| 10–50 kb – pairs | 513 | 829 |
| 10–50 kb – % of total | 9.74% | 15.70% |
| 10–50 kb – bases (bp) | 10,331,331 | 16,129,629 |
| 50–100 kb – pairs | 104 | 139 |
| 50–100 kb – % of total | 1.97% | 2.63% |
| 50–100 kb – bases (bp) | 7,556,688 | 9,567,717 |
| >100 kb – pairs | 55 | 143 |
| >100 kb – % of total | 1.04% | 2.71% |
| >100 kb – bases (bp) | 10,956,637 | 41,504,577 |

[1] Number of duplicated segment pairs (each pair = two genomic regions that are copies of each other). Filtered to >90% identity, >=1 kb avg size, chromosomes only.

[2] Unique genomic bases inside at least one SD region. Overlapping SD regions are merged so each base is counted once.

[3] Sum of (size1+size2)/2 across all pairs WITHOUT merging overlaps. Overcounts hotspot regions where multiple SDs overlap. Use merged coverage instead for manuscript.

[4] SD pairs where both copies are on the same chromosome.

[5] Non-redundant genomic bases in intra-chromosomal SD regions.

[6] Sum of avg pair sizes for intra-chromosomal SDs, without merging.

[7] SD pairs where copies are on different chromosomes.

[8] Non-redundant genomic bases in inter-chromosomal SD regions.

[9] Sum of avg pair sizes for inter-chromosomal SDs, without merging.

**Table S13. Gene-impacting variants distinguishing prairie and meadow vole genomes, related to Figure 3.**

| Metric | Value |
|---|---|
| **Ortholog groups (OrthoFinder)** | |
| Total ortholog groups | 31,831 |
| 1:1 orthologs | 17,571 |
| Expanded in meadow | 539 |
| Expanded in prairie | 533 |
| Multi-copy in both | 78 |
| Meadow-specific | 7,040 |
| Prairie-specific | 6,070 |
| **Coding-region variants (autosomal CDS, prairie reference)** | |
| Total CDS variants | 415,006 |
| Synonymous | 259,445 (62.5%) |
| Missense | 142,712 (34.4%) |
| Stop gained | 807 (0.2%) |
| Stop lost | 153 (0.0%) |
| Start lost | 263 (0.1%) |
| Splice acceptor | 165 (0.0%) |
| Splice donor | 222 (0.1%) |
| Splice region | 5,185 (1.2%) |
| Frameshift insertion | 727 (0.2%) |
| Frameshift deletion | 747 (0.2%) |
| In-frame insertion | 2,090 (0.5%) |
| In-frame deletion | 1,939 (0.5%) |
| HIGH-impact projections (VEP) | 3,135 variants (2,166 genes) |
| **Candidate LGD genes filtered (1:1 ortholog)** | |
| VEP candidate LGD | 839 variants (295 genes) |
| SnpEff candidate LGD | 1,043 variants (261 genes) |
| Cross-annotator consensus genes | 253 |
| VEP projections in consensus | 755 |
| SnpEff projections in consensus | 1,027 |
| **Selection pressure (Ka/Ks, Nei–Gojobori 1986)** | |
| Genome-wide 1:1 orthologs with Ka/Ks | 17,610 |
| Median Ka/Ks | 0.111 |
| Mean Ka/Ks | 0.179 |
| Genes with Ka/Ks ≥ 1 | 233 |
| LoF consensus with Ka/Ks | 250 / 253 |
| Median Ka/Ks | 0.292 |
| Mean Ka/Ks | 0.361 |
| Genes with Ka/Ks ≥ 1 | 5 |

#### Supplemental Methods

**Methods S1. Comparison of CiFi vs Hi-C in genome assembly, scaffolding, and haplotype phasing performance, related to Figure 1.** CiFi-phased contigs were comparable or longer in three of four haplotypes and consolidated into fewer final scaffolds, approaching expected chromosome numbers more closely than Hi-C (Figures S2 and S3, Table S2). For example, prairie vole Hap1 yielded 63 scaffolds with CiFi versus 171 with Hi-C (Figure S3). Hi-C also accumulated 1.7–2× more scaffold joins across YAHS rounds yet produced more final scaffolds, and introduced approximately twice the gap sequence across all four assemblies (Figure S4), suggesting that many Hi-C joins did not persist in the final assembly. Where Hi-C showed a higher scaffold N50 in the meadow vole, this was accompanied by more gaps. Overall, CiFi achieved better or equivalent scaffold consolidation, supporting its utility as a single-platform approach for chromosome-resolved diploid *de novo* assembly.

We next downsampled CiFi reads from 100% (5× coverage) to 1% in each species to determine the minimum input required to generate contiguous genomes (Figure S5). Assembly sizes remained stable across all conditions and haplotypes, while scaffold N50 and counts were both sensitive to reduced CiFi depth. All four haplotypes maintained near-optimal scaffold N50 at 40% input or 5 Gbp of CiFi data (~2× genome coverage). Extending this test to a human genome (NA12878) shows highly accurate haplotype phasing: median per-chromosome switch error was 0.18% versus 0.38% for HiFi only (30×), and essentially equal to standard 30× Hi-C (0.16%) (Figure S6A, Table S3). Increasing CiFi to 5× further reduced switch error by 39% (0.11%), with continued monotonic improvement at higher depths, to 0.04% at the full SMRT Cell CiFi yield (~24×).

When considering the Hamming error, the HiFi-only (30×) baseline has a low median per-chromosome switch error rate (0.38%) but a median blockwise Hamming error rate of 47% (Figure S6B, Table S3). Thus, local phase relationships between adjacent heterozygous variants are often correct, but phase consistency does not extend across the chromosome. Adding 2× CiFi reduces the median Hamming error rate to 0.45%, a more than a 100-fold reduction, and full-depth CiFi (24×) reduces it to 0.05%. CiFi also has a lower Hamming error rate than Hi-C at every shared downsampled depth (for example, 0.45% versus 1.33% at 2×); at the maximum available depth of each library, CiFi reaches 0.05% at 24× versus 0.31% for Hi-C at 30×. These results demonstrate that a single CiFi library supports accurate chromosome-scale haplotype phasing, a conclusion that the switch-error analysis alone could not establish.
